# A Framework for Transcriptome-Wide Association Studies in Breast Cancer in Diverse Study Populations

**DOI:** 10.1101/769570

**Authors:** Arjun Bhattacharya, Montserrat García-Closas, Andrew F. Olshan, Charles M. Perou, Melissa A. Troester, Michael I. Love

**Affiliations:** Department of Biostatistics, University of North Carolina at Chapel Hill; Division of Cancer Epidemiology and Genetics, National Cancer Institute; Division of Genetics and Epidemiology, Institute of Cancer Research, London, UK; Department of Epidemiology, University of North Carolina at Chapel Hill; Lineberger Comprehensive Cancer Center, University of North Carolina at Chapel Hill; Department of Genetics, University of North Carolina at Chapel Hill; Department of Pathology and Laboratory Medicine, University of North Carolina at Chapel Hill

**Keywords:** transcriptome-wide analysis (TWAS), breast cancer, expression quantitative trait loci (eQTL), survival, polygenic traits

## Abstract

**Background:** The relationship between germline genetic variation and breast cancer survival is largely unknown, especially in understudied minority populations who often have poorer survival. Genome-wide association studies (GWAS) have interrogated breast cancer survival but often are underpowered due to subtype heterogeneity and many clinical covariates and detect loci in non-coding regions that are difficult to interpret. Transcriptome-wide association studies (TWAS) show increased power in detecting functionally-relevant loci by leveraging expression quantitative trait loci (eQTLs) from external reference panels in relevant tissues. However, ancestry- or race-specific reference panels may be needed to draw correct inference in ancestrally-diverse cohorts. Such panels for breast cancer are lacking.

**Results:** We provide a framework for TWAS for breast cancer in diverse populations, using data from the Carolina Breast Cancer Study (CBCS), a North Carolina population-based cohort that oversampled black women. We perform eQTL analysis for 406 breast cancer-related genes to train race-stratified predictive models of tumor expression from germline genotypes. Using these models, we impute expression in independent data from CBCS and TCGA, accounting for sampling variability in assessing performance. These models are not applicable across race, and their predictive performance varies across tumor subtype. Within CBCS (*N = 3,828*), at a false discovery-adjusted significance of 0.10 and stratifying for race, we identify associations in black women near *AURKA, CAPN13, PIK3CA, and SERPINB5* via TWAS that are underpowered in GWAS.

**Conclusions:** We show that carefully implemented and thoroughly validated TWAS is an efficient approach for understanding the genetics underpinning breast cancer outcomes in diverse populations.

## Background

Breast cancer remains the most common cancer among women in the world [1]. Breast cancer tends to be more aggressive in young women and African American women, though underlying germline determinants of poor outcomes are not well-studied. Cohorts that represent understudied minority populations, like the Carolina Breast Cancer Study (CBCS), have identified differences in healthcare access, socioeconomics, and environmental exposures associated with disparities in outcome [2–4], but more targeted genomic studies are necessary to interrogate these disparities from a biologic and genetic perspective.

Few genome-wide association studies (GWAS) have studied the relationship between germline variation and survival outcomes in breast cancer, with most focusing instead on genetic predictors of risk [5, 6]. Recently, GWAS have shown evidence of association between candidate common germline variants and breast cancer survival, but these studies are often underpowered [7, 8]. Furthermore, the most significant germline variants identified by GWAS, in either risk or survival, are often located in non-coding regions of the genome, requiring *in vitro* follow-up experiments and co-localization analyses to interpret functionally [9]. It is important to seek strategies for overcoming these challenges in GWAS, especially because several studies in complex traits and breast cancer risk have shown that regulatory variants not significant in GWAS account for a large proportion of trait heritability [10–12].

Novel methodologic approaches that integrate multiple data types offer advantages in interpretability and statistical efficiency. Escala-García et al. has suggested that aggregating variants by integrating gene expression or other omics may better explain underlying biological mechanisms while increasing the power of association studies beyond GWAS [7]. To alleviate problems with statistical power and interpretability, a recent trend in large-scale association studies is the transcriptome-wide association study (TWAS). TWAS aggregates genomic information into functionally-relevant units that map to genes and their expression. This gene-based approach combines the effects of many regulatory variants into a single testing unit that increases study power and provides more interpretable trait-associated genomic loci [13–15]. Hoffman et al. and Wu et al. have recently conducted TWAS for breast cancer risk and have reported several significant associations for genes with breast cancer susceptibility, showing increased power over GWAS [15, 16]. However, these studies either draw from ancestrally-homogeneous reference panels like subsets of women of European ancestry from the Genotype-Tissue Expression (GTEx) project [16] or study populations of European descent from the Breast Cancer Association Consortium (BCAC) [15]. It is not known whether these models can be informative in African American women and other groups. Recent findings have suggested that stratification by race or ancestry may be necessary to construct proper tests of association across race or ancestry [17, 18]. However, many cohorts, especially large-scale genetic cohorts, may not have a sufficient sample size in minority populations to power these tests.

Here, we provide a framework for TWAS for complex disease outcomes in diverse study populations using transcriptomic reference data from the Carolina Breast Cancer Study (CBCS), a multi-phase cohort that includes an over-representation of African American women [19]. We train race-stratified predictive models of tumor expression from germline variation and carefully validate their performance, accounting for sampling variability and disease heterogeneity, two aspects that previous TWAS in breast cancer have not considered. This framework shows promise for scaling up into larger GWAS cohorts for further detection of risk- or outcome-associated loci.

## Results

### Race specific germline eQTL analysis

To assess the association between germline genomic variation and tumor expression of 406 autosomal genes, targeted by the CBCS because of their association with breast cancer progression, we first conducted a full cis-trans expression quantitative trait loci (eQTL) analysis, stratifying on race and controlling for key biological covariates and population stratification (see **Methods**). We discuss the relationship between self-reported race and ancestry in CBCS in **Supplemental Results**.

We evaluated associations between the tumor expression levels of 406 autosomal genes and 5,989,134 germline SNPs. SNPs and genes found in association in an eQTL will be called eSNPs and eGenes, respectively. At a Benjamini-Bogomolov [20] FDR-corrected *P*-value (*BBFDR < 0.05*), we identified 266 cis-eQTLs and 77 trans-eQTLs in the AA sample across 32 eGenes, and 691 cis-eQTLs and 15 trans-eQTLs in the WW sample across 24 eGenes, shown in **Supplemental Figure 2**. Of these eGenes, 4 are in common across race: *PSPHL*, *GSTT2*, *EFHD1*, and *SLC16A3*. Expressions of *PSPHL* and *GSTT2* have been previously reported to be governed by respective cis-deletions and serve as distinguishing biomarkers for race [21–24]. The majority of significant eQTLs in both the AA and WW samples were found in cis-association with respective eGenes. However, we saw a higher proportion of significant trans-eQTLs in the AA sample (**Supplemental Figure 2**). The locations and strengths of top eQTLs for all 406 autosomal genes are shown in Figure 1A. All significant eQTLs are plotted in **Supplemental Figure 2**.

**Figure 1:**
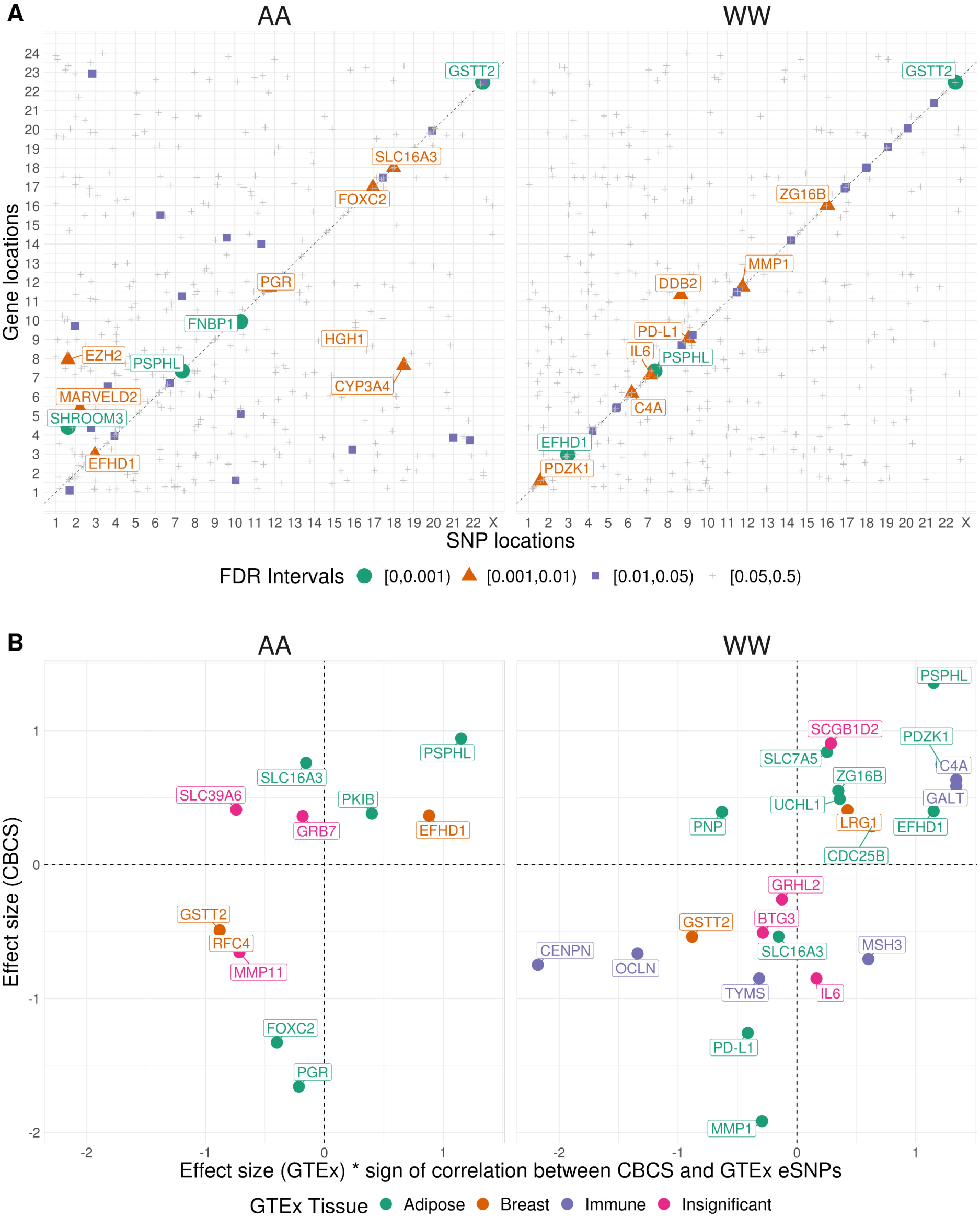
(A) Cis-trans plot of top eQTL by gene stratified by SRR. Each point represents the top eQTL for a given gene. The color and size of each point reflects the Benjamini-Bogomolov FDR-adjusted P-value (BBFDR) for that eQTL. eGenes with *BBFDR < 0.01* are labelled. (B) Comparison of effect sizes of eGenes with significant cis-eQTLs in CBCS (Y -axis) and GTEx (X-axis) over tissue type, stratified by race. eGenes are colored by the GTEx tissue that shows the largest effect size. GTEx effect sizes on the X-axis are multiplied by the sign of the correlation between the genotypes of the GTEx and CBCS eSNPs.

We further adjusted our eQTL models for a computationally-derived estimate of tumor purity, which showed little effect on the strength and location of top eQTLs by eGene (**Supplemental Results**). We do not consider tumor purity in any downstream analyses and train predictive models on bulk tumor expression.

We lastly sought to evaluate the source of the significant eQTLs we detect in CBCS. Similar to previous pan-cancer germline eQTL analyses [25], we cross-referenced eGenes found in CBCS with eGenes detected in relevant healthy tissues from Genotype-Tissue Expression (GTEx) Project. We attributed all but 7 of the cis-eGenes from CBCS across both AA and WW women found in GTEx to one of these three tissue types (Figure 1B), with the effect sizes of the top eQTLs for these eGenes correlating very well between CBCS and GTEx (see **Supplemental Figure 5**).

### Race-specific predictive models of tumor expression

Using the significant germline eQTLs of tumor expression as motivation, we used tumor expression and genotyping data from 628 AA women and 571 WW women from CBCS to build predictive models of tumor RNA expression levels for each gene’s breast tumor expression (see **Methods**). Mean cis-heritability (cis-*ℎ^2^*) of the 417 genes is 0.016 (*SE = 0.019*) in AA women and 0.015 (*SE = 0.019*), as estimated by GREML-LDMS analysis [26]. For downstream analysis, we only consider genes with cis-*ℎ^2^* significantly greater than 0 at a nominal *P*-value less than 0.10 from the relevant likelihood ratio test. Considering only these genes, the mean cis-*ℎ^2^* of genes is 0.049 (*SE = 0.016*) in AA models and 0.052 (*SE = 0.016*) in WW models. Of the predictive models built for these genes, 125 showed a five-fold cross-validation prediction performance (CV *R^2^*) of at least 0.01 (10% Pearson correlation between predicted and observed expression with *P < 0.05*) in one of the two predictive models. Figure 2A shows the CV *R^2^*of these 153 genes across race. The median CV *R^2^*for the 153 genes was 0.011 in both AA and WW women. Cis-*ℎ^2^* and CV *R^2^* are compared in **Supplemental Figure 6**.

**Figure 2:**
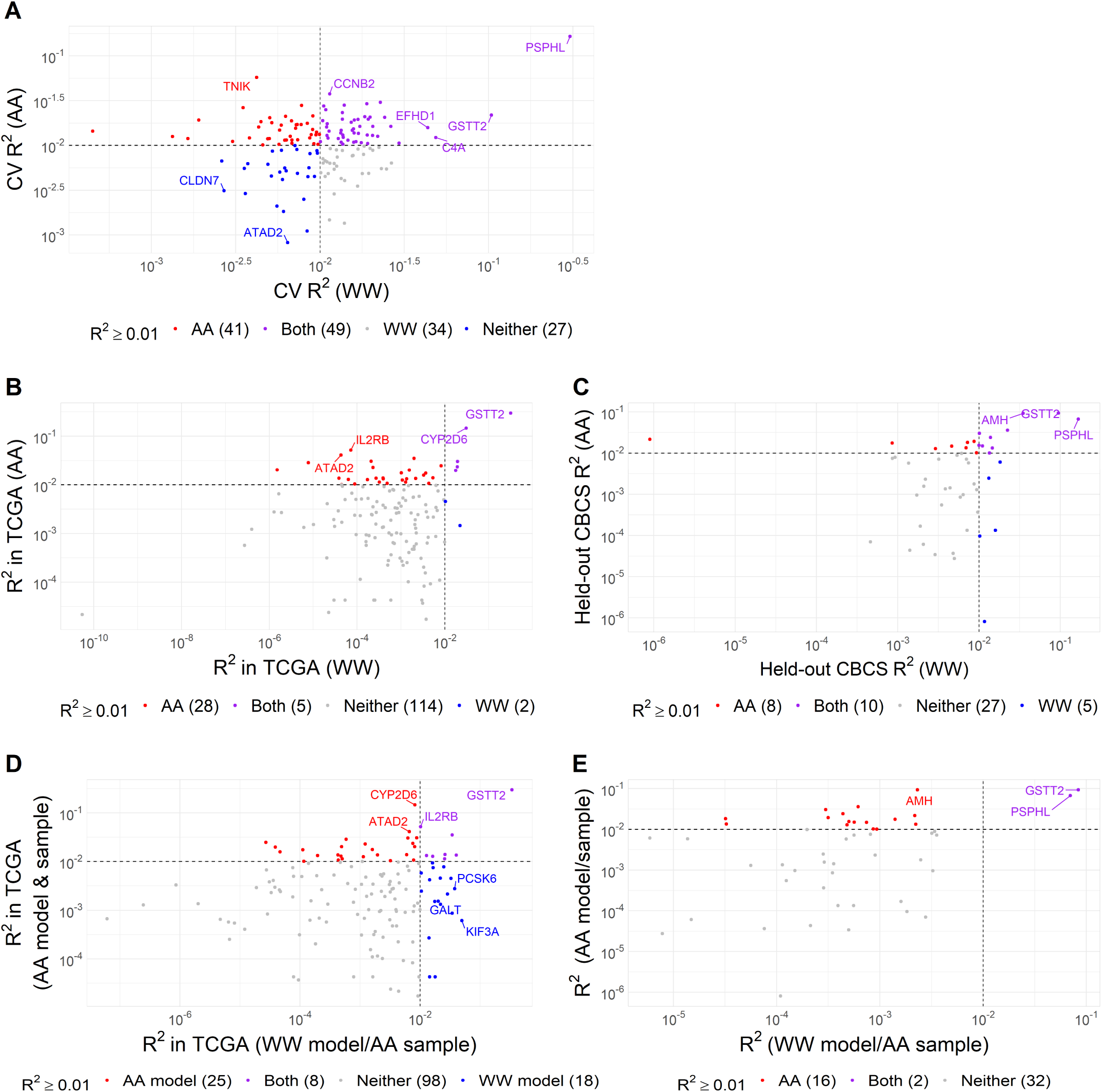
(A) Comparison of cross-validation *R^2^* across race in CBCS. Cross-validation *R^2^* in CBCS WW women (X-axis) and CBCS AA women (Y-axis) for each of the 151 analyzed genes. Scales are logarithmic. Dotted lines represent *R^2^ = 0.01*. Colors represent the model with which a given gene can be predicted at *R^2^ > 0.01*. (B) Cross-validation *R^2^* in CBCS (X-axis) and square Spearman correlation between observed expression and GReX in TCGA-BRCA (Y-axis) in AA sample (left) and WW sample (right). Pearson correlations between *R^2^* calculated on the raw scale. are plotted on the log-scale. (C) Comparison of validation *R^2^*across race in TCGA for 149 analyzed genes found in TCGA expression data. (D) Comparison

Based on model performance in CBCS, we selected 46 genes in AA women and 57 genes in WW women for association analyses between predicted tumor gene expression and breast cancer survival, using data from all patients from CBCS with genotype data. These genes were selected because they showed an CV *R^2^ > 0.01* (10% correlation between observed and predicted expression in the CBCS training set) and cis-*ℎ^2^ ≥ 0* with nominal *P < 0.10* in a given race strata.

### Evaluation of predictive models in independent data

Predictive performance was strong across race and biological and molecular subtype in two external samples: The Cancer Genome Atlas (TCGA) and a held-out CBCS sample set. We defined the imputed expression of a given gene in an external cohort as the GReX, or the germline-genetically regulated tumor expression, of that gene.

The first sample is derived from TCGA breast tumor tissues with 179 AA and 735 WW women. We compared predictive performance by calculating an external validation *R^2^* (EV *R^2^*) with squared Spearman correlations. Of the 151 genes modeled in CBCS training data with significant cis-*ℎ^2^*, 149 genes were measured via RNA-seq in TCGA. A comparison of predictive performance in TCGA for these 149 genes is shown in Figure 2B, showing adequate performance in AA women (33 genes with EV *R^2^> 0.01*) and poor performance in WW women (7 genes with EV *R^2^> 0.01*). The top predicted gene in cross-validation from CBCS for both races, *PSPHL*, was not present in the TCGA normalized expression data and could not be validated. Another top cross-validated gene, *GSTT2*, was present in TCGA expression data and was validated as the top genetically predicted gene in TCGA by EV *R^2^*.

We also imputed expression into entirely held-out samples from CBCS data (1,121 AA and 1,070 WW women) that have gene expression for a subset of the genes (166 of 417 genes) in the CBCS training set. These samples were largely derived from Phases I and II of CBCS (see **Methods**). A comparison of imputation performance in CBCS for 51 genes is shown in Figure 2C, showing adequate performance in both AA and WW women (18 and 15 genes with EV *R^2^ > 0.01* in AA and WW women).

### Predictive models are not applicable across race

We find that the predictive accuracy of most genes was lower when expression was imputed in AA women using models trained in the WW sample. We employed the WW predictive models to impute expression into AA samples from TCGA and held-out CBCS data. We compare the performances of the WW model and AA model in the AA sample in Figure 2D (TCGA) and **2E** (CBCS). In held-out CBCS samples, with the WW model, we could only predict *PSPHL* and *GSTT2* at *R^2^ > 0.01* in the AA sample, as the expression of these genes is modulated mostly by strongly associated cis-eSNPs. In TCGA, our WW models performed adequately in AA women, though the WW models predicted fewer genes at *R^2^ > 0.01* than the AA models.

### Evaluation of predictive performance across subtype

While predictive accuracy of expression models was stable across datasets, there was greater heterogeneity across biological and molecular subtype. In part, this is due to small sample sizes within race and subtype-specific strata. Upon first inspection, we see vast differences in the performance of our models across subtype (**Supplemental Figure 7**), with a large majority of genes performing at EV *R^2^ > 0.01* in rarer subtypes, like HER2-enriched breast cancers. However, we recognized sample sizes in the TCGA validation set were relatively small, especially when considering AA women and women of certain subtype, e.g. as low as 16 AA women with HER2-enriched breast cancer. As overall correlation between observed and imputed expressions are near 0, we sought to account for sampling variability when imputing into groups of women with such small sample sizes.

We employed a permutation scheme: permuting observed expression values among samples 10,000 times to generate a null distribution for EV *R^2^*. We then tested for the null hypothesis *R^2^ = 0*, controlling for false discovery, according to this null distribution. Supplemental Figure 9 displays *q*-values in Manhattan form [27], showing that the proportion of genes with EV *R^2^* significantly different from 0 is similar across subtypes. We inverted this permutation test [28] to construct a confidence interval for EV *R^2^*. We find that the EV *R^2^* of several genes are highly variable across subtypes, even when accounting for differences in sample size and therefore sampling variation. Key examples of such genes with variable EV *R^2^* across subtypes are shown in Figure 3.

**Figure 3:**
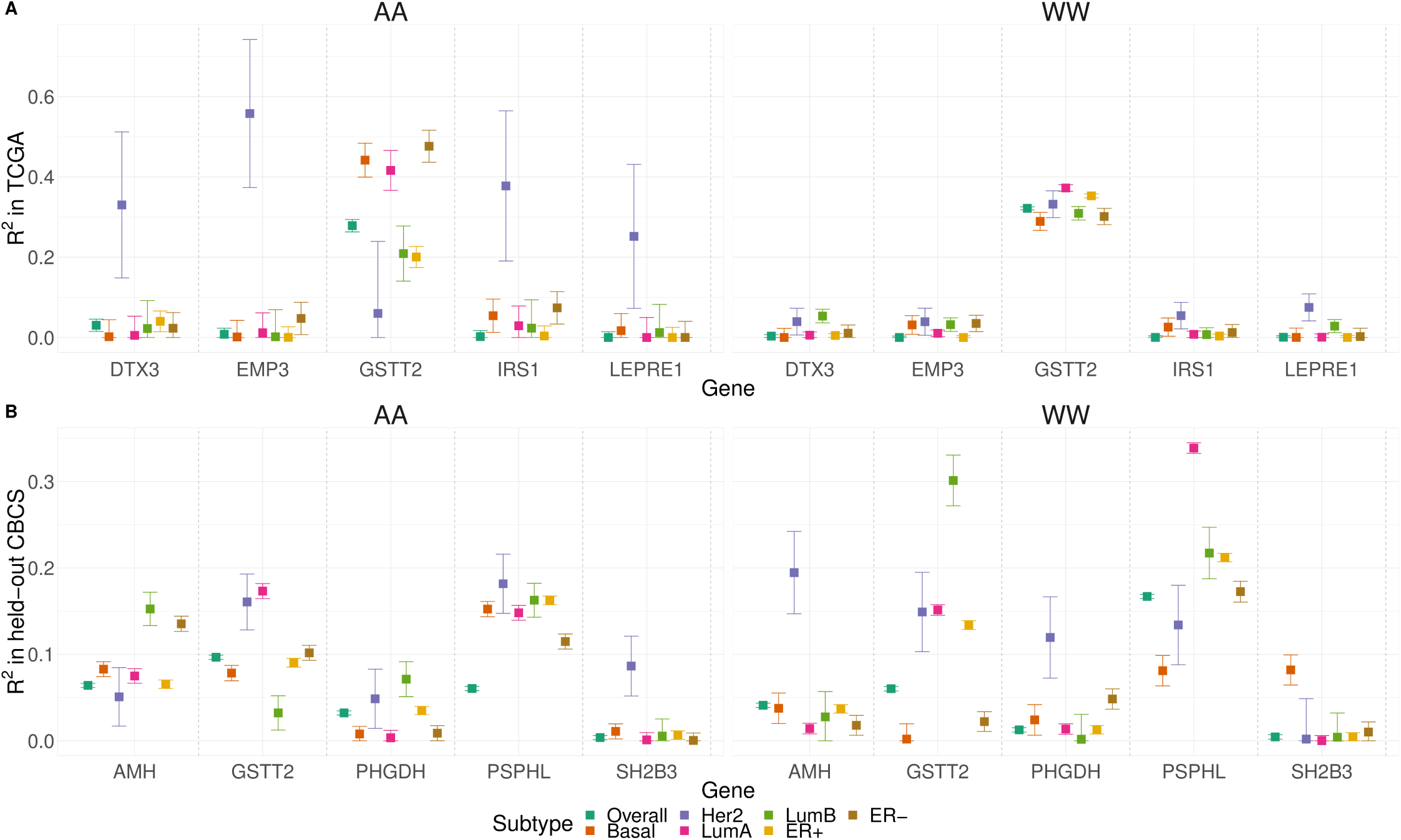
Validation *R*^2^ across PAM50 molecular subtype and estrogen receptor status, stratified by race, for example genes with highly variable *R*^2^ in TCGA (A) and held-out CBCS (B). Squared Spearman correlation (Y-axis), denoted *R*^2^, between observed and predicted gene expression is plotted for different genes (X-axis), stratfied by PAM50 subtype and estrogen receptor status. Points are colored and shaped according to subtype. Error bars provide 90% confidence intervals inverted from the corresponding permutation test.

### Predicted expression associated with breast cancer-specific survival

To assess association between imputed gene expression and breast cancer-specific survival, we constructed race-stratified cause-specific proportional hazard models for 3,828 samples from CBCS (1,865 AA and 1,963 WW), where we model time to mortality due to breast cancer. Of the genes evaluated, we detected 4 whose GReX were associated with breast-cancer specific survival at FDR-adjusted *P < 0.10* in AA women, shown in Table 1 and Figure 4. We did not identify any genes with GReX associated with survival in WW women.

**Figure 4:**
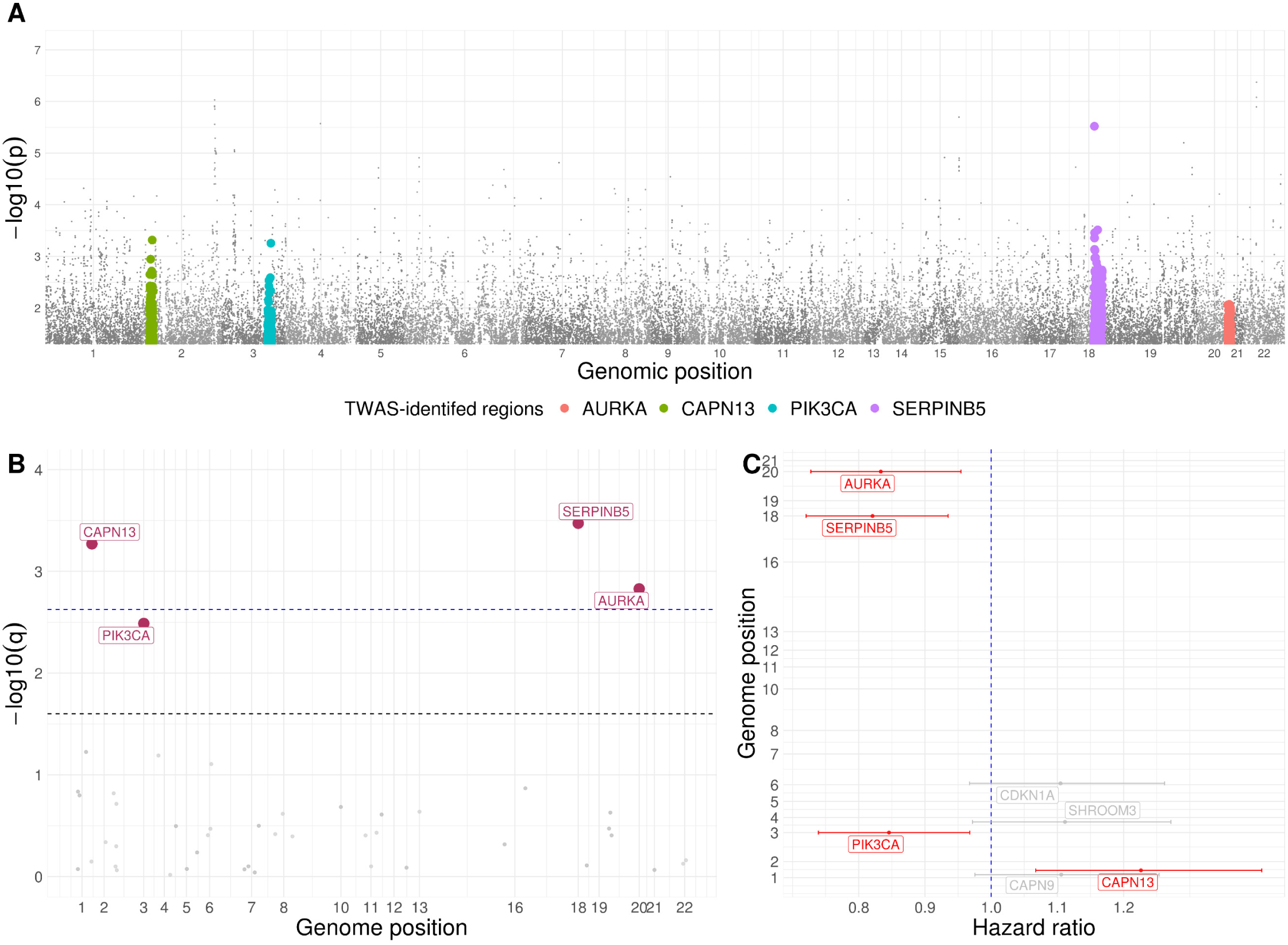
GWAS and TWAS results in AA women. (A) Manhattan plot of traditional GWAS on breast cancer survival. Genomic regions found to be significantly associated with survival in TWAS are represented in various colors. No SNVs reach Benjamini-Hochberg FDR-adjusted genome-wide significance. (B) Manhattan plot of TWAS on breast cancer survival. Genomic regions found to be significant at FDR-adjusted *P < 0.10* are highlighted in red. The blue line represents a cutoff of FDR-adjusted *a = 0.05* and the dotted black line represents a cutoff of FDR-adjusted *a = 0.10*. (C) Caterpillar plot of log-hazard rates with FDR-adjusted 90% confidence levels (X-axis) and genomic position (Y-axis). Results shown are significant at nominal *P < 0.10*. Genes highlighted in red represent genes with GReX significantly associated with survival at FDR-adjusted *P < 0.10*.

**Table 1:** Genes with GReX found in association with breast cancer-specific survival in AA women. (a) Hazard ratio and FDR-adjusted 90% confidence intervals, *Z*-statistic, and *P*-value of association of GReX with breast cancer-specific survival. (b) Cross-validation *R^2^* of gene expression in AA models.

| Region | Gene | Hazard Ratio<br>(90% CI)<sup>a</sup> | Z-Statistic<sup>a</sup> | P-value<sup>a</sup> | GReX $R^2$<sup>b</sup> |
| --- | --- | --- | --- | --- | --- |
| 20q13.2 | AURKA | 0.83 (0.73, 0.95) | -2.52 | $1.5 \times 10^{-3}$ | 0.021 |
| 2p23.1 | CAPN13 | 1.22 (1.07, 1.41) | 2.76 | $5.4 \times 10^{-4}$ | 0.011 |
| 3q26.32 | PIK3CA | 0.85 (0.74, 0.97) | -2.34 | $3.2 \times 10^{-3}$ | 0.013 |
| 18q21.33 | SERPINB5 | 0.82 (0.72, 0.93) | -2.85 | $3.4 \times 10^{-4}$ | 0.010 |

An association between increased GReX and increased risk of breast cancer-specific mortality was identified for *CAPN13* (*2p23.1*). We also found protective associations between higher GReX of *AURKA (20q13.2)*, *PIK3CA (3q26.32)*, *SERPINB5* (*18q21.33*) and lower risk of breast cancer-mortality (Figure 4C). Of these 4 loci, associations with survival have been reported with SNPs in the same chromosomal region as *AURKA*, *PIK3CA,* and *SERPINB5* [8,29–33], though none of these reported SNPs were utilized in constructing the GReX of this gene. Furthermore, the GReX of these four genes were not significantly correlated (*P > 0.05* for all pairwise Spearman correlation tests), and the sets of SNPs used in constructing the GReX of these four genes had no pairwise intersections, providing evidence that their independent association with breast cancer-specific survival was not a pleiotropic effect from shared or correlated SNPs.

To determine whether the associations between predicted gene expression and breast cancer-specific survival were independent of GWAS-identified association signals, we performed conditional analyses adjusted for the most significant GWAS-identified survival-associated SNPs closest to the TWAS-identified gene by adjusting the cause-specific proportional hazards model for the genotype from this SNP. We found that the association for *PIK3CA* had a small change in effect size after adjustment for its adjacent survival-associated SNP, and its SNP-adjusted association was insignificant, while the other genes’ associations remained significant after adjustment (Table 2). This conditional analysis suggests that the GReX of *AURKA*, *CAPN13*, and *SERPINB5* may be associated with breast cancer-specific survival independent of the GWAS-identified variant. No previously reported survival-associated SNPs were found significant at the genome-wide significance level in our dataset, and none of the closest survival-associated SNPs used in conditional adjustment were significant (Figure 4A). This supports our observation that correctly analyzed TWAS using relevant tissue gene expression may increase power for association testing.

**Table 2:** Genes with GReX found in association with breast cancer-specific survival. (a) Top survival-associated SNP in cis-region of the given gene from GWAS for survival and distance of top cis-SNP from gene. (b) FDR-adjusted hazard ratio, 90% confidence interval, and *P*-value for association of GReX and breast cancer-specific survival, adjusting for adjacent survival-associated SNPs.

| Gene | Closest survival-associated SNP<sup>a</sup> | Distance to closest survival-associated SNP<sup>a</sup> | Hazard ratio, adjusting for adjacent GWAS-SNP (90% CI)<sup>b</sup> | P-value, adjusting for adjacent risk SNPs<sup>b</sup> |
| --- | --- | --- | --- | --- |
| AURKA | rs202100873 | 87.1 kb | 0.84 (0.74, 0.94) | 0.027 |
| CAPN13 | rs72068647 | 266.9 kb | 1.18 (1.04, 1.33) | 0.046 |
| PIK3CA | rs66487567 | 271.9 kb | 0.88 (0.78, 1.00) | 0.096 |
| SERPINB5 | rs376302305 | 89.4 kb | 0.84 (0.75, 0.94) | 0.028 |

As we deal with case-only data, we wished to inspect any collider bias that arises from unmeasured confounders that are associated with both breast cancer incidence and survival (see **Supplemental Figure 1**3) [34]. Since a case-control dataset was not readily available to us to test associations between the GReX of genes with breast cancer risk, we construct the weighted burden test, as in FUSION [14], for the GReX of *AURKA*, *CAPN13, PIK3CA*, and *SERPINB5* in the GWAS summary statistics for breast cancer risk in AA women available from BCAC using the iCOGs dataset and additional GWAS [35–37]. We find that none of the GReX of these genes are significantly associated with breast cancer incidence (*Z > 1.96, P < 0.05*), suggesting minimal presence of collider bias in our estimates of association with survival for the GReX of these four genes.

Lastly, we examined the association of the GReX of these four genes with breast cancer-specific survival in AA women, stratified by estrogen receptor (ER) subtype. We find that overall associations with survival are often driven by significant associations in a single subtype, though there is evidence of significant hazardous association in both ER subtypes for *CAPN13* (**Supplementary Figure 1**0). We also did not detect a survival association with the total expression of these 4 genes, as estimated from breast cancer-specific Cox models (**Supplementary Figure 11**).

## Discussion

In this paper, we studied the relationship between breast cancer-specific survival and germline genetics using a TWAS framework, wherein we aggregate the germline genome into testing units that map to the transcriptome to greatly mitigate the multiple testing burden found in GWAS. This study is the first systematic TWAS for breast cancer-specific survival, motivated by a full cis-trans eQTL analysis with one of the largest sample sizes for breast tumor gene expression in African American women. Our analyses underscore the importance of accounting for sampling variability when validating predictive models for TWAS and incorporating race or ancestry in these models, an aspect which confounds naïve comparisons involving imputed GReX across validation sub-groups of different sample size.

Using a training set from CBCS, we leveraged race-stratified germline eQTLs of tumor expression to train race-stratified models of tumor expression from germline variation. Our eQTL analysis reveals a strong cis-signal between germline variants and tumor expression of several genes, that is both differential across race and not exclusively attributable to healthy breast tissue. Our models showed strong cross-validation predictive performance in genes with significant cis-heritability. We also show strong predictive performance in a held-out test set from CBCS and adequate performance of our WW models in TCGA-BRCA data. We suspect that this discrepancy in validation performance between CBCS and TCGA may be attributed to a poor intersection of SNPs in the genotyping data from TCGA and CBCS (only approximately 85% of SNPs from CBCS represented in TCGA imputed genotype data). There could also be a lack of cis-heritability of the tumor expression of a majority of genes assayed in TCGA. For example, Gusev et al. has trained models for gene expression in breast tumors in TCGA; only 8 of the 417 genes in the CBCS Nanostring panel showed significant cis-heritability in their models [14], which we downloaded from the Gusev Lab’s TWAS/FUSION repository. We believe that predictive performance in TCGA data consistent with CBCS data is a high bar for validation due to both genotyping and RNA expression platform differences between CBCS (Oncoarray and Nanostring) and TCGA (Affymetrix 6.0 and RNAseq). Reproducible performance in both AA and WW women in our independent test set from CBCS data suggests that our models are quite robust. Follow-up studies, in which models of tumor expression are trained in TCGA RNA-seq data and validated in CBCS Nanostring data, could elucidate any discrepancies in predictive performance across platform.

An important implication of our work is the race-specificity of TWAS methods. In our validation scheme, we assessed the applicability of imputing expression in AA samples using the WW predictive models, as publicly available tumor expression data is often measured in predominantly WW cohorts. We find that WW models generally have poor performance in AA women. Epidemiological studies have stressed accounting for differences in race by stratification or adjustment for admixture estimates when constructing polygenic scores [38]. Our key finding of poor predictive performance across race suggests that this epidemiological note of caution extends to creating predictive models for RNA expression. Previous TWAS studies of breast cancer risk have either used models trained in a sample of predominantly European ancestries [16] or imputed into large cohorts of strictly patients of European descent [15]. Hoffman et al. does exclude SNPs that were monomorphic in any of the 14 different ancestral populations they analyze [16], though this may not capture all effects of ancestry on genetic regulation of expression, including the possibility for interactions. We contend that accounting for ancestry or stratifying by race may be necessary to draw correct inference in large, ancestrally-heterogeneous cohorts.

Our data also suggests that predictive performance may vary by molecular subtype. Previous groups have shown the predictive utility of catering polygenic risk scores to breast cancer subtype [39, 40], a phenomenon we investigated in our predictive models of tumor expression. As the estimates of sample correlations between observed and predicted expression were small and the sample sizes per subtype were small, we recognized the need to employ a permutation method to assess the precision of our prediction *R^2^*. We found that a significant portion of the variability of predictive performance across subtype was explained by sampling variability. Nevertheless, even after accounting for sampling variability, we noticed that several genes have varied predictive performance across subtype and race. This finding suggests that TWAS predictive models of expression may need to account of biological heterogeneity. We also reinforce the importance of sampling variability in the validation of predictive models in external cohorts prior to generalized imputation and association testing. For example, Wu et al. trained their models in a relatively small set of 67 women from GTEx and validated their 12,824 models in a validation set of 86 women from TCGA without accounting for sampling variability of predictive performance [15]. A recent multi-tissue TWAS in ovarian cancer from Gusev et al. considered a more thorough validation of predictive models by leveraging multiple independent cohorts to assess replication rates for their models [41]. We recommend such an approach if multiple independent cohorts are accessible. But, in TWAS evaluation in a single tissue, studies should place a strong emphasis on validation, accounting for sampling variability of prediction *R^2^*, ideally prior to imputation in larger cohorts.

While many of the most significant findings here are methodological in nature, we also have data to suggest that four genomic loci may merit further investigation relative to breast cancer survival. We identified 4 genomic loci associated with breast cancer survival at an FDR-adjusted significance level of 0.10 in AA women. After adjustment for genetics at the most significantly survival-associated SNP close to the gene in question, survival associations at 3 of these 4 locations remained marginally significant. We did not observe any significant association between the total expression of these 4 genes and breast cancer-specific survival. This suggests that the germline-regulated component of the tumor expression of these genes – a small fraction of the total expression variation – may be associated with survival outcomes. Numerous factors, including copy number alterations, epigenetic or post-transcriptional regulation, and exposures and technical artifacts in measurement contributed to the total expression measured in the tumor. Thus, we do not expect that significant GReX association implies total expression association, or vice versa.

While nearly all of the genes on the CBCS Nanostring panel are relevant to breast cancer research, many have not been shown to be associated with survival. Two of these 4 TWAS-identified genes have strong functional evidence in breast cancer survival literature. Mutations in *AURKA* and *PIK3CA* have previously been shown to be significantly associated with breast cancer survival rates [29–31]. Less is known about the involvement of *SERPINB5* and *CAPN13* in breast cancer survival. *SERPINB5* is a tumor-suppressor gene that has been shown to promote development of breast cancers in humans [42]. The calpain family, which contain *CAPN13*, is a group of proteases that is involved in apoptosis and the progression and proliferation of breast cancer cells and has been suggested as therapy targets for various cancers [43–45]. These four loci merit further studies for validation and functional characterization, both in large GWAS cohorts and using *in vitro* studies.

We also observed that 3 of the 4 associations were driven by very strong effect sizes within a single subtype (Supplementary Figure 11). Though we cannot contextualize this result, it highlights an often-overlooked modeling consideration. In a cohort that is both biologically and ancestrally-heterogenous, as in CBCS, investigators should consider modeling choices beyond simple linear adjustments for subtype and race. Given a large enough sample size, it may be prudent in future TWAS to stratify predictive models on both race and biological subtype to increase power to detect outcome-associated loci that are strongly present within only one such strata or have heterogeneous effects across strata. This idea is akin to the logic of Begg et al and Martínez et al in detecting etiological risk factors for ER-positive and -negative tumors [46, 47].

Since the CBCS analysis was a case-only study, we were wary of potential collider bias by unmeasured confounders associated with both breast cancer risk and progression [34,48–50]. These colliders may affect the magnitude and direction of effect sizes on association between survival and GReX of genes (Supplemental Figure 14). We find that, using summary statistics for breast cancer risk GWAS from iCOGs [35–37], none of the GReX of these four genes showed significant transcriptome-wide associations with breast cancer risk in this iCOGs data. This suggests that our estimates of association may be free of the collider bias, outlined in Supplemental Figure 14. As Escala-García et al. highlights, germline variation can affect breast cancer prognosis via tumor etiology (risk of developing a tumor of a certain subtype), or via mechanisms that are relevant post-tumorigenesis, such as the cellular response to therapy, or the host-tumor micro-environment, including immune response and stroma-tumor interactions [7]. Ideally, in future TWAS and integrated omic analyses of breast cancer survival, it is prudent to consider joint models of breast cancer risk and survival to account for the many effects of germline genotype and any associations with unmeasurable confounders [49].

One limitation of our study is that data on somatic amplifications and deletions were not yet available for the CBCS cohort we analyzed. Removing the somatic copy number variation signal from tumor expression profiles may improve our estimates of cis-heritability and perhaps the predictive performance of our models, as previous TWAS have shown [41]. Furthermore, not all genes in the CBCS Nanostring panel have a significant heritable component in expression regulation. These genes, like *ESR1*, which have a significant role in breast cancer etiology [51], could not be investigated in our study. Lastly, since CBCS mRNA expression is assayed by the Nanostring nCounter system, we could only analyze 94 aggregated locations on the human transcriptome across race. However, the Nanostring platform allows the CBCS to robustly measure expression from FFPE samples on a targeted panel of breast cancer and race-related genes, allowing us to leverage the large sample size from all three phases of the CBCS. One of the greatest strengths of our study is that the CBCS affords us both a large training and test set of AA and WW women for race-stratified predictive models. Such data is important in drawing inference in more ancestrally-heterogeneous populations. Accordingly, the statistical power of our study is high to detect associations for genes with relatively high cis-heritability. Nonetheless, the specific survival-associated loci merit further investigation in external datasets. Future studies in large GWAS cohorts, such as those within the Breast Cancer Association Consortium, will elucidate how to account for ancestral and biological heterogeneity in detecting survival-associated loci.

## Conclusion

We have provided a framework of transcriptome-wide association studies (TWAS) for breast cancer outcomes in diverse study populations, considering both ancestral and subtype-dependent biological heterogeneity in our predictive models. From a more theoretical perspective, this work will inform the utilization of TWAS methods in polygenic traits and diverse study populations, stressing rigorous validation of predictive models prior to imputation and careful modeling to capture associations with outcomes of interest in diverse populations.

## Methods

### Data collection

#### Study population

The Carolina Breast Cancer Study (CBCS) is a population-based study conducted in North Carolina (NC) that began in 1993; study details and sampling schemes have been described in previous CBCS work [19, 52]. Patients of breast cancer aged between 20 and 74 years were identified using rapid case ascertainment in cooperation with the NC Central Cancer Registry, with self-identified African American and young women (ages 20-49) oversampled using randomized recruitment [19]. Randomized recruitment allows sample weighting to make inferences about the frequency of subtype in the NC source population. Details regarding patient recruitment and clinical data collections are described in Troester et al [2].

Date of death and cause of death were identified by linkage to the National Death Index. All diagnosed with breast cancer have been followed for vital status from diagnosis until date of death or date of last contact. Breast cancer-related deaths were classified as those that listed breast cancer (International Statistical Classification of Disease codes 174.9 and C-50.9) as the underlying cause of death on the death certificate. By the end of follow-up, we identified 674 deaths, 348 of which were due to breast cancer. In total, we compiled 3,828 samples (1,865 AA and 1,963 WW) from all phases of CBCS with relevant survival and clinical variables.

#### CBCS genotype data

Approximately 50% of the SNPs for the OncoArray were selected as a “GWAS backbone” (Illumina HumanCore), which aimed to provide high coverage for the majority of common variants through imputation. The remaining SNPs were selected from lists supplied by six disease-based consortia, together with a seventh list of SNPs of interest to multiple disease-focused groups. Approximately 72,000 SNPs were selected specifically for their relevance to breast cancer. The sources for the SNPs included in this backbone, as well as backbone manufacturing, calling, and quality control, are discussed in depth by the OncoArray Consortium [53]. All samples were imputed using the October 2014 (v.3) release of the 1000 Genomes Project dataset as a reference panel in the standard two-stage imputation approach, using *SHAPEIT2* for phasing and *IMPUTEv2* for imputation [54–56]. All genotyping, genotype calling, quality control, and imputation was done at the DCEG Cancer Genomics Research Laboratory [53].

From the provided genotype data, we excluded variants (1) with a minor frequency less than 5% and (2) that deviated significantly from Hardy-Weinberg equilibrium at *P < 10^-8^* using the appropriate functions in *PLINK v1.90b3* [57, 58]. Finally, we intersected genotyping panels for the AA and WW samples, resulting in 5,989,134 autosomal variants and 334,391 variants of the X chromosome. CBCS genotype data was coded as dosages, with reference and alternative allele coding as in the National Center for Biotechnology Information’s Single Nucleotide Polymorphism Database (dbSNP).

#### CBCS gene expression data

Paraffin-embedded tumor blocks were requested from participating pathology laboratories for each sample, reviewed, and assayed for gene expression using Nanostring as discussed previously [2]. In total, 1,388 samples with invasive breast cancer from the CBCS were analyzed for a total of 406 autosomal genes and 11 genes on the X chromosome. All assays were performed in the Translational Genomics Laboratory at the University of North Carolina at Chapel Hill.

We used the *NanoStringQCPro* package in Bioconductor to first eliminate samples that did not have sufficient Nanostring data quality [59]. Next, we normalized distributional differences between lanes with upper-quartile normalization [60]. Unwanted technical and biological variation (i.e. tissue heterogeneity) was estimated in the resulting gene expression data with techniques from the *RUVSeq* package from Bioconductor [61]. Unwanted variation was controlled using the distribution of 11 endogenous housekeeping genes on the Nanostring gene expression panel. Ultimately, we removed 2 dimensions of unwanted variation from the variance-stabilized transformation of the gene expression data [62, 63]. We lastly used principal component analysis to detect and remove any significant, potential outliers. A final intersection of samples that had both genotype and gene expression data gave us a final sample of 1,199 subjects (628 AA women and 571 WW women).

#### TCGA genotype data

Birdseed genotype files of 914 of WW and AA women were downloaded from the Genome Data Commons (GDC) legacy (GRCh37/hg19) archive. Genotype files were merged into a single binary PLINK file format (BED/FAM/BIM) and imputed using the October 2014 (v.3) release of the 1000 Genomes Project dataset as a reference panel in the standard two-stage imputation approach, using SHAPEIT v2.837 for phasing and IMPUTE v2.3.2 for imputation [54–56]. We excluded variants (1) with a minor allele frequency of less than 1%, (2) that deviated significantly from Hardy-Weinberg equilibrium (*P < 10^-8^*) using appropriate functions in PLINK v1.90b3 [57, 58], and (3) located on sex chromosomes. We further excluded any SNPs not found on the final, quality-controlled CBCS genotype data. Final TCGA genotype data was coded as dosages, with reference and alternative allele coding as in dbSNP.

#### TCGA expression data

TCGA level-3 normalized RNA expression data were downloaded from the Broad Institute’s GDAC Firehose (2016/1/28 analysis archive) and subsetted to the 417 genes analyzed in CBCS. A total of 412 of these 417 were available in TCGA expression data.

### Computational methods

#### Deconvolution of bulk tumor RNA

A study pathologist analyzed tumor microarrays (TMAs) from 176 of the 1,199 subjects to estimate area of dissections originating from epithelial tumor, assumed here as a proxy for the proportion of the bulk RNA expression attributed to the tumor. Using these 176 observations as a training set and the normalized gene expressions as the design matrix, we trained a support vector machine model tuned over a 10-fold cross-validation [64, 65]. The cross-validated model was then used to estimate tumor purities for the remaining 1,023 samples from their gene expressions. We do not consider tumor purity in final eQTL models and all downstream analyses.

#### eQTL analysis

We assessed the additive relationship between the gene expression values and genotypes with linear regression analysis using *MatrixeQTL* [66], in the following model:

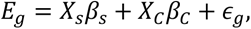

where *Ẽ_g_* is the gene expression of gene *g*, *X_s_* is the vector of genotype dosages for a given SNP *s*, *C* is a matrix of covariates, *β_s_*and *β_C_*are the effect-sizes on gene expression for the SNP *s* and the covariates *C*, respectively, and *E* is assumed to be Gaussian random error with mean 0 and common variance *a^2^* for all genes *g*.

We calculated both cis-(variant-gene distance less than 500 kb) and trans-associations between variants and genes. Classical *P*-values were calculated for Wald-type tests of *H_o_: β_s_ = 0* and were adjusted post-hoc via the Benjamini-Bogomolov hierarchical error control procedure, *TreeQTL* [20]. We conducted all eQTL analyses stratified by race. Age, BMI, postmenopausal status, and the first 5 principal components of the joint AA and WW genotype matrix were included in the models as covariates in *C*. Estimated tumor purity was also included as a covariate to assess its impact on strength and location of eQTLs. Any SNP found in an eQTL with Benajmini-Bogomolov adjust *P*-value *BBFDR < 0.05* is defined as an eSNP using *TreeQTL* [20]. The corresponding gene in that eQTL is defined as an eGene. We exclude samples with Normal-like subtype, as classified by the PAM50 classifier, due to generally low tumor content.

We downloaded healthy tissue eQTLs from the Genotype-Tissue Expression (GTEx) Project and cross-referenced eGenes and corresponding eSNPs between CBCS and GTEx in healthy breast mammary tissue, EBV-transformed lymphocytes, and subcutaneous adipose tissue. The Genotype-Tissue Expression (GTEx) Project was supported by the Common Fund of the Office of the Director of the National Institutes of Health, and by NCI, NHGRI, NHLBI, NIDA, NIMH, and NINDS. The data used for the analyses described in this manuscript were obtained from the GTEx Portal on 05/12/19.

#### Estimation of cis-heritability

Cis-heritability (cis-*ℎ^2^*) was estimated using the GREML-LDMS method, proposed to estimate heritability by correction for bias in linkage disequilibrium (LD) in estimated SNP-based heritability [26]. Analysis was conducted using *GCTA* v.1.92 [67]. For downstream analysis, we only consider the 151 genes (81 in AA women and 100 in WW women) with cis-*ℎ^2^* that can be estimated with nominal *P*-value *< 0.10*.

#### Predictive tumor expression models

We adopt general techniques from PrediXcan and FUSION to estimate eQTL-effect sizes for predictive models of tumor expression from germline variants [13, 14]. First, gene expressions were residualized for the covariates *C* included in the eQTL models (age, BMI, postmenopausal status, and genotype PCs) given the following ordinary least squares model:

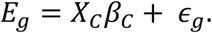

We then consider downstream analysis on 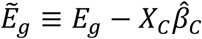

For a given gene *g*, we consider the following linear predictive model:

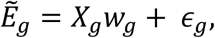

where *E^-^_g_*is the gene expression of gene *g*, residualized for the covariate matrix *X_C_*, *X_g_*is the genotype matrix for gene *g* that includes all cis-SNPs for gene *g* (within 500 kb of either the 5’ or 3’ end of the gene) and all trans-eQTLs with *BBFDR < 0.01*, *w_g_* is a vector of effect-sizes for eQTLs in *X_g_*, and *E_g_* is Gaussian random error with mean 0 and common variance for all *g*.

We estimate *w_g_* with the best predictive of three schemes: (1) elastic-net regularized regression with mixing parameter *a = 0.5* and *;i_* penalty parameter tuned over 5-fold cross-validation [13, 68], (2) linear mixed modeling where the genotype matrix *X_g_* is treated as a matrix of random effects and *ŵ_g_*is taken as the best linear unbiased predictor (BLUP) of *w_g_*, using *rrBLUP* [69], and (3) multivariate linear mixed modeling as described above, estimated using *GEMMA* v.0.97 [70].

In these models, the genotype matrix *X_g_* is pruned for linkage disequilibrium (LD) prior to modeling using a window size of 50, step size of 5, and LD threshold of 0.5 using *PLINK* v.1.90b3 [58] to account for redundancy in signal. The final vectors *ŵ_g_*of effect-sizes for each gene *g* are estimated by the estimation scheme with the best 5-fold cross-validation performance. All predicted models are stratified by race, i.e. an individual model of tumor expression for AA women and WW women for each gene *g*.

To impute expression into external cohorts, we then construct the germline genetically-regulated tumor expression *GReX_g_*of gene *g* given *ŵ_g_*in the predictive model as follows:

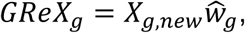

where *X_g,new_*is the genotype matrix of all available SNPs in the feature set of *ŵ_g_*in a GWAS cohort.

All final models are available here: https://github.com/bhattacharya-a-bt/CBCS_TWAS_Paper.

#### Validation in TCGA

Using our stratified predictive models of tumor expression, we imputed expression in TCGA and measured predictive accuracy of each gene through prediction *R^2^*, defined here as the squared Spearman correlation between observed and imputed expression. It is important to note that all variants in the CBCS-trained predictive models are not represented in the TCGA genotype data. Predictive performance in TCGA was also assessed stratified by PAM50 intrinsic subtype and estrogen receptor status.

To account for sampling variability in calculating correlations in validation cohorts of smaller sample sizes, we calculated a permutation null distribution for each gene by permuting observed expressions 10,000 times and calculating a “null” prediction *R^2^* at each permutation. The sample validation prediction *R^2^*was compared to this permutation null distribution to generate an empirical *P*-value for the sample *R^2^*, using Storey’s *qvalue* package. We then calculated *q*-values from these empirical *P*-values, controlling for a false discovery rate of 0.05 [27]. Lastly, we constructed confidence intervals for *R^2^* by inverting the acceptance region from the permutation test [28].

#### Validation in CBCS

We used an entirely held-out sample of 2,308 women from CBCS as a validation set of Nanostring nCounter data on a codeset of 166 genes. These samples were normalized as outlined before. We used the same validation methods as in TCGA, as well using a permutation method to assess the statistical significance of predictive performance, stratified by PAM50 subtype and estrogen receptor status.

#### PAM50 subtyping

GReX in CBCS were first estimated as outlined above. We residualized the original tumor expression *E* for these imputed expression values to form a matrix of tumor expression adjusted for GReX (*E^-^*). We then classified each subject into PAM50 subtypes based on both *E* and *E^-^*, using the procedure summarized by Parker et al [71, 72].

#### Survival modeling

Here, we defined a relevant event as a death due to breast cancer. We aggregated all deaths not due to breast cancer as a competing risk. Any subjects lost to follow-up were treated as right-censored observations. We estimated the association of GReX with breast cancer survival by modeling the race-stratified cause-specific hazard function of breast cancer-specific mortality, stratifying on race [73]. For a given gene *g*, the model has form

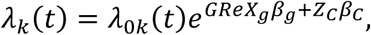

where *β_g_*is the effect size of *GReX_g_*on the hazard of breast cancer-specific mortality, *Z_C_* represents the matrix of covariates (age at diagnosis, estrogen-receptor status at diagnosis, tumor stage at diagnosis, and study phase), and *β_C_* are the effect sizes of these covariates on survival. *λ_k_(t)* is the hazard function specific to breast cancer mortality, and *λ_ok_(t)* is the baseline hazard function. We test *H_o_: β_g_= 0* for each gene *g* with Wald-type tests, as in a traditional Cox proportional hazards model. We correct for genomic inflation and bias using *bacon*, a method that constructs an empirical null distribution using a Gibbs sampling algorithm by fitting a three-component normal mixture on *Z*-statistics from TWAS tests of association [74].

Here, we consider only the 46 genes that have CV *R^2^ > 0.01* in AA women and the 57 genes that have CV *R^2^ > 0.01* in WW women for race-stratified survival modeling. We adjust tests for *β_g_*via the Benjamini-Hochberg procedure at a false discovery rate of 0.10.

For comparison, we run a GWAS to analyze the association between germline SNPs and breast cancer-specific survival using *GWASTools* [75]. We use a similar cause-specific hazards model with the same covariates as in the TWAS models of association, correcting for false discovery with the Benjamini-Hochberg procedure.

#### Inspection of collider bias

To assess collider bias when conditioning for breast cancer incidence in case-only studies, such as CBCS, we test for association for the GReX of genes with breast cancer risk using iCOGs summary statistics from BCAC [35–37], using the weighted burden test identified by FUSION [14]. In summary, we compose a weighted *Z* test statistic as follows:

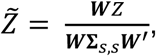

where *Z* is the vector of *Z*-statistics from iCOGs and 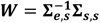 with *∑_e,s_* is the covariance matrix between all SNPs represented in *Z* and the gene expression of the given gene and *I_s,s_* is the covariance among all SNPs.

#### Power analysis

Using *survSNP* [76], we generated the empirical power of a GWAS to detect various hazard ratios with 3,828 samples with 1,000 simulation replicates at a significance level of *P = 1.70 × 10^-8^*, corresponding to an FDR-adjusted *P = 0.10*. We assume an event rate of 10%, a relative allelic frequency of the risk allele of 0.1 and estimate the 90^th^ percentile of times-to-event as a landmark time. Similarly, for genes of various cis-*ℎ^2^*, we assessed the power of TWAS to detect various hazard ratios at *P = 0.0096* (corresponding to FDR-adjusted *P = 0.10*) over 1,000 simulation replications from the empirical distribution function of the GReX of the given gene.

## Supporting information

Supplemental Results

## Abbreviations

CBCS: Carolina Breast Cancer Study
GWAS: Genome-wide association study
LD: Linkage disequilibrium
SNP/V: Single nucleotide polymorphism/variant
TWAS: Transcriptome-wide association study
GTEx: The Genoype-Tissue Expression Project
BCAC: Breast Cancer Association Consortium
PRS: Polygenic risk score
WW: self-identified white women
AA: self-identified African American women
ER: estrogen receptor
eQTL: expression quantitative trait loci
AMBER: Alberta Moving Beyond Breast Cancer
eGene: eQTL-associated gene
eSNP: SNP found in an eQTL
FDR: false discovery rate
BBFDR: Benjamini-Bogomolov adjusted false discovery rate
*ℎ^2^*: heritability
TCGA: The Cancer Genome Atlas
BRCA: breast cancer
GReX: germline-genetically regulated tumor expression

## Declarations

### Ethics approval and consent to participate

This study was approved by the Office of Human Research Ethics at the University of North Carolina at Chapel Hill, and written informed consent was obtained from each participant.

### Consent for publication

Not applicable

### Availability of data and materials

Summary statistics eQTL results, tumor expression models, and relevant R code for training expression models in CBCS are freely available at https://github.com/bhattacharya-a-bt/CBCS_TWAS_Paper/.

### Competing interests

C.M.P. is an equity stockholder in and consultant for BioClassifier LLC; C.M.P. is also listed as an inventor on patent applications on the Breast PAM50 Subtyping assay. The other authors declare that they have no competing interests.

### Funding

Susan G. Komen® provided financial support for CBCS study infrastructure. Funding was provided by the National Institutes of Health, National Cancer Institute P01-CA151135, P50-CA05822, and U01-CA179715 to A.F.O, C.M.P. and M.A.T. M.I.L. is supported by R01-HG009937, R01-MH118349, P01-CA142538, and P30-ES010126.

The Translational Genomics Laboratory is supported in part by grants from the National Cancer Institute (3P30CA016086) and the University of North Carolina at Chapel Hill University Cancer Research Fund. Genotyping was done at the DCEG Cancer Genomics Research Laboratory using funds from the NCI Intramural Research Program.

The funders had no role in the design of the study, the collection, analysis, or interpretation of the data, the writing of the manuscript, or the decision to submit the manuscript for publication.

### Authors’ contributions

A.B., M.G., A.F.O., M.A.T., and M.I.L. conceived the study. A.B. developed the statistical approaches, performed the analysis, and drafted the paper. A.B., M.A.T., and M.I.L. performed initial exploratory analysis. C.M.P., M.A.T., and M.I.L. provided insight in methodological approaches and analysis. M.G., A.F.O., C.M.P., and M.A.T. provided data resources. M.A.T. and M.I.L. supervised the study. All authors approved and edited the final manuscript.

## Acknowledgements

We thank the Carolina Breast Cancer Study participants and volunteers. We also thank Colin Begg, Jianwei Cai, Nilanjan Chatterjee, Alexander Gusev, Katherine Hoadley, Yun Li, John Witte, and Emily Zabor for valuable discussion during the research process. We thank Erin Kirk and Jessica Tse for their invaluable support during the research process. We thank the DCEG Cancer Genomics Research Laboratory and acknowledge the support from Stephen Chanock, Rose Yang, Meredith Yeager, Belynda Hicks, and Bin Zhu.

## Notes

https://github.com/bhattacharya-a-bt/CBCS_TWAS_Paper

