## Supplemental Results for "A Framework for Transcriptome-Wide Association Studies in Breast Cancer in Diverse Study Populations"

*Admixture and genotype principal components correlate with self-reported race*

In downstream analyses (eQTL analysis and training of predictive tumor expression), we stratify on self-reported race. We understand that self-reported race is not fully reflective of genetic ancestry. However, **Supplemental Figure 1** shows that self-reported race correlates strongly with population stratification in the CBCS. Furthermore, we see a linear relationship between African admixture estimates and the first genotype principal component of representative samples from the CBCS cohort that are included in the Alberta Moving Beyond Breast Cancer (AMBER) Cohort Study [1]. For all downstream eQTL analysis and tests of associations, we control for the first five principal components of the combined self-reported African American (AA) and self-reported white (WW) genotype matrix, accounting for differences in ancestry prior to stratification by self-reported race. We refer to self-reported race as race throughout this work.

*Tumor purity adjustment and overlap with healthy tissue eQTLs*

Geeleher et al. show that only a third of conventional eQTLs in bulk breast cancer tumor expression could be attributed to cancer cells in TCGA [2]. We wished to assess the extent to which this observation bore out in CBCS. For this reason, we developed an estimate of tumor purity to use as an adjustment covariate for our eQTL analysis (see **Methods**). In general, we do not see significant differences in the strength and location of significant eQTLs, as shown in comparative cis-trans plots of all eQTLs across race and adjustment for tumor purity (**Supplemental Figures 3 and 4**). For most genes, top eQTL regions with linkage disequilibrium (LD) support had small differences in strength of effect size (Manhattan plot for a representative gene shown in Supplemental Figure 3). Adjusting for tumor purity, at $\text{BBFDR} < 0.05$, we identified 266 cis-eQTLs and 84 trans-eQTLs in the AA sample across 36 eGenes, and 634 cis-eQTLs and 14 trans-eQTLs in the WW sample across 23 eGenes, shown in Supplemental Figure 4. All WW eGenes, adjusting for tumor purity, are in common with WW eGenes from bulk tumor expression, and 32 of 36 AA eGenes, adjusting for tumor purity, are in common with AA eGenes from bulk tumor expression. Top eQTLs for eGenes remain largely the same across adjustment for tumor purity. Summary statistics for these eQTLs are provided, as mentioned in **Availability of data and materials**. Due to limited differences when we adjust for tumor purity, all downstream analyses do not involve our computational estimate of tumor purity.

We do not observe the same difference in eQTLs across adjustment for tumor purity as in Geeleher et al [2]. The Nanostring expression data from CBCS includes only 417 genes, all of which were selected for the panel because of their involvement in breast cancer tumorigenesis, biology, or outcome disparities due to race. Furthermore, our normalization procedure involves the RUV method, which accounts for unwanted technical and biological variation, estimated from the distributions of housekeeping negative controls [3,4] with an unsupervised method. We hypothesize that the RUV method accounts for a significant percentage of the variability from cell-type heterogeneity that may confound traditional eQTL analysis in bulk tumor RNA expression. Further implementations of deconvolution algorithms specialized for expression measured for targeted panels of genes, as in Nanostring, would aid in distinguishing the source cell types or tissues for various breast tumor eQTLs. Accurate bulk expression deconvolution may also be important in future TWAS to consider sources of variation in tumor expression due to tissue heterogeneity and how deconvoluted tumor expression signals contribute to outcomes of interest.

*PAM50 subtype calls robust to adjustment for GReX*

Emami et al. has shown that genetically regulated tumor expression can elucidate biological mechanisms that can delineate prostate cancer subtypes [6]. Using tumor expression data in held-out CBCS, we assessed differences in PAM50 molecular subtype calls before and after adjustment by imputed GReX. We adjusted bulk TCGA tumor expression by imputed GReX and ran PAM50 subtyping on full and GReX-adjusted tumor expression [7]. Of the 2,174 samples analyzed for PAM50 subtyping, only 15 show different PAM50 subtypes after adjusting for GReX (**Supplementary Figure 9A**). Most of these discordant pairs (12 out of 15) are between the HER2-enriched, Luminal A, and Luminal B subtypes, all of which are relatively similar in terms of proliferation and their molecular profiles. Discordant pairs show relatively similar confidence scores (defined as $1 - P$-value of Spearman correlation to PAM50 subtype centroid), proliferation scores, and ROR-P scores [7], shown in **Supplementary Figures 10B-D**. The robustness of PAM50 subtype calling to GReX adjustment is consistent with our observation that only 10 of the 50 genes comprising the PAM50 gene signature were cis-heritable at $P < 0.10$ in our data set.

*Power in TWAS to detect survival associations*

Previous studies have suggested increased power in TWAS to detect smaller effect sizes in studies of disease risk [8,9]. We generated the empirical power of a GWAS to detect various hazard ratios with 3,828 samples using 1,000 simulation replicates with an event rate, risk allele frequency, landmark time and probabilities to landmark times derived from CBCS genotype data at a significance level of $P = 1.70 \times 10^{-8}$, corresponding to a FDR-adjusted $P = 0.10$ [10]. Similarly, for simulated genes with various cis-$h^2$, we assessed the power of a hypothetical TWAS analysis to detect various gene-mediated hazard ratios at $P = 0.0096$ (corresponding to FDR-adjusted $P = 0.10$) over 1,000 simulation replications from the empirical distribution function of the GReX. It is important to note that the detectable hazard ratios at 80% for GWAS and TWAS are incomparable due to differences in units of measure. At 80% power, a GWAS with CBCS data with $N = 3{,}828$ is powered to detect a hazard ratio of breast cancer-specific survival of 1.88 with an addition of one alternative allele in a given SNP. At 80% power, in our study, TWAS can detect hazard ratios 1.186, 1.203, and 1.216 with the GReX of a gene with cis-$h^2 \approx$ 0.100, 0.055, and 0.030, with respect to an increase of one standard deviation, respectively (**Supplemental Figure 12**).
