## Supplemental Figures for "A Framework for Transcriptome-Wide Association Studies in Breast Cancer in Diverse Study Populations"

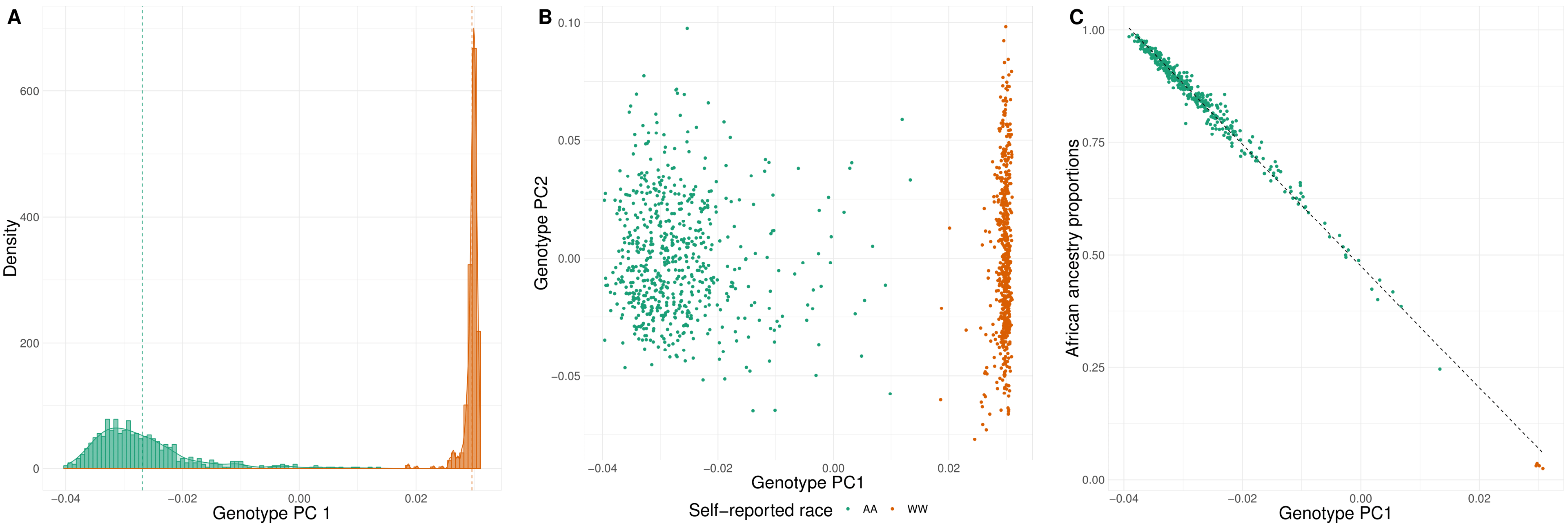


**Figure 1**: (A) Density plot of first principal component of genotype matrix, colored by self-reported race. (B) PCA plot of first principal component of genotype matrix (X-axis) and second principal component of genotype matrix (Y-axis), colored by self-reported race. (C) Plot of genotype PC1 (X-axis) against African admixture ancestry estimates from the Alberta Moving Beyond Breast Cancer (AMBER) Study Cohort (Y-axis). The sample plotted here is an intersection of patients from CBCS and the AMBER cohort.

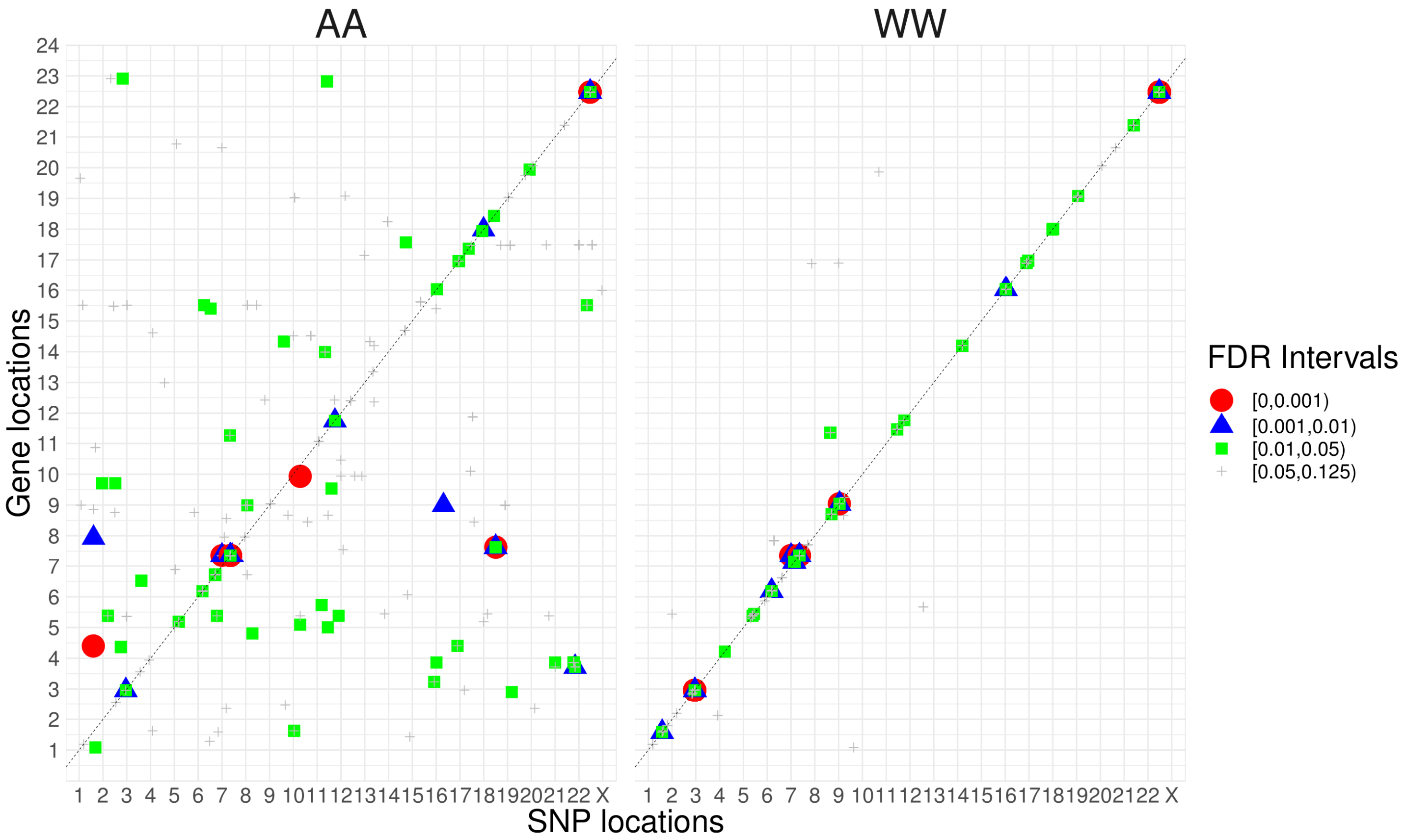


**Figure 2**: Cis-trans plot of race-stratified eQTL analyses, AA on the left and WW on the right. Each point represents an eQTL with $BBFDR < 0.125$ with the location of the 5' end of the corresponding eGenes on the X-axis and the genomic location of the corresponding eSNP on the Y-axis. A 45-degree line is provided as a reference for cis-eQTLs.

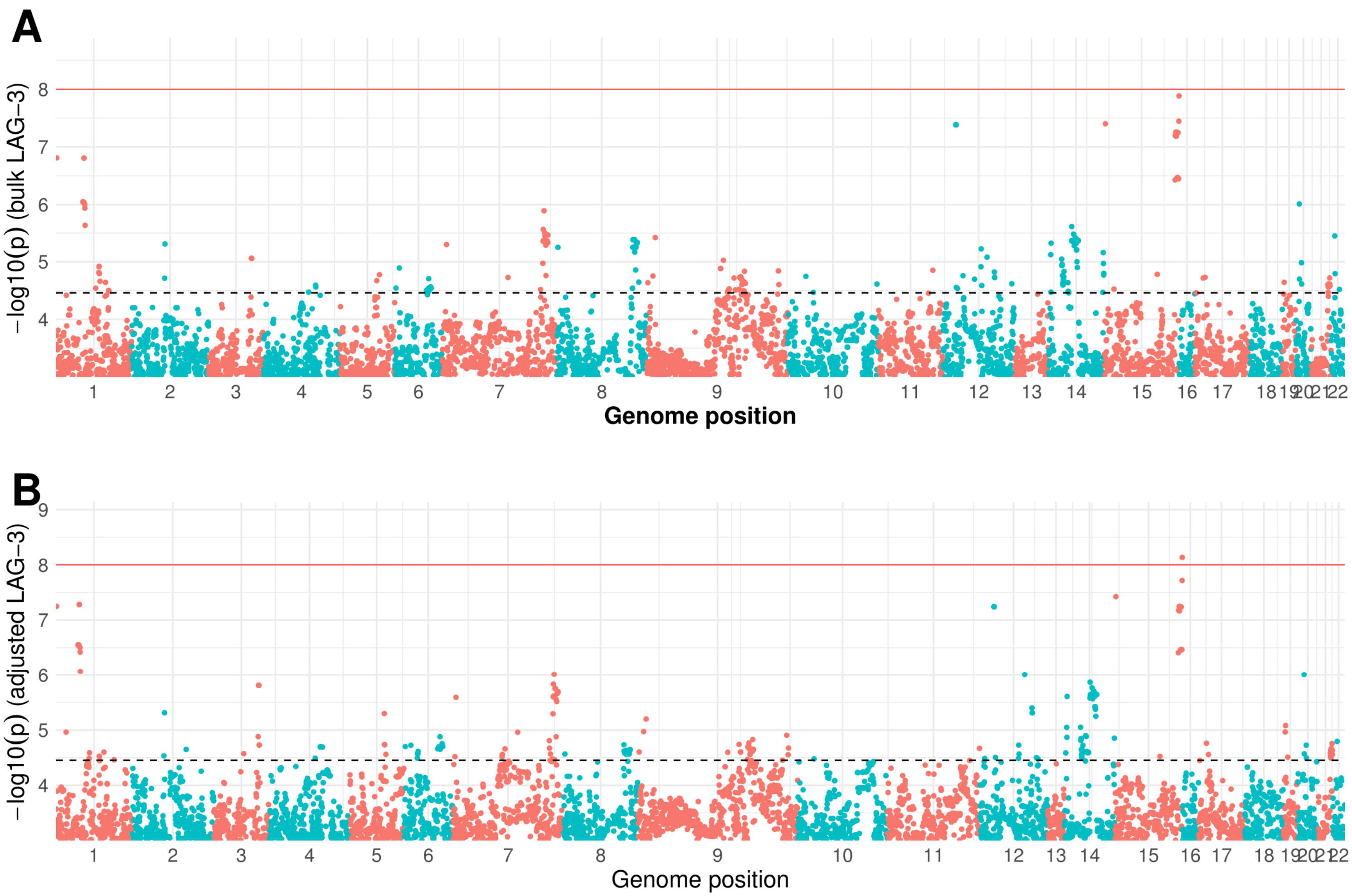


**Figure 3**: Example Manhattan plots for eQTL analysis in bulk tumor *LAG-3* expression (A) and tumor purity-adjusted *LAG-3* expression (B) in WW women. Red line represents a genome-wide significance threshold of $P = 1 \times 10^{-8}$ and the dotted black line corresponds to $BBFDR < 0.05$.

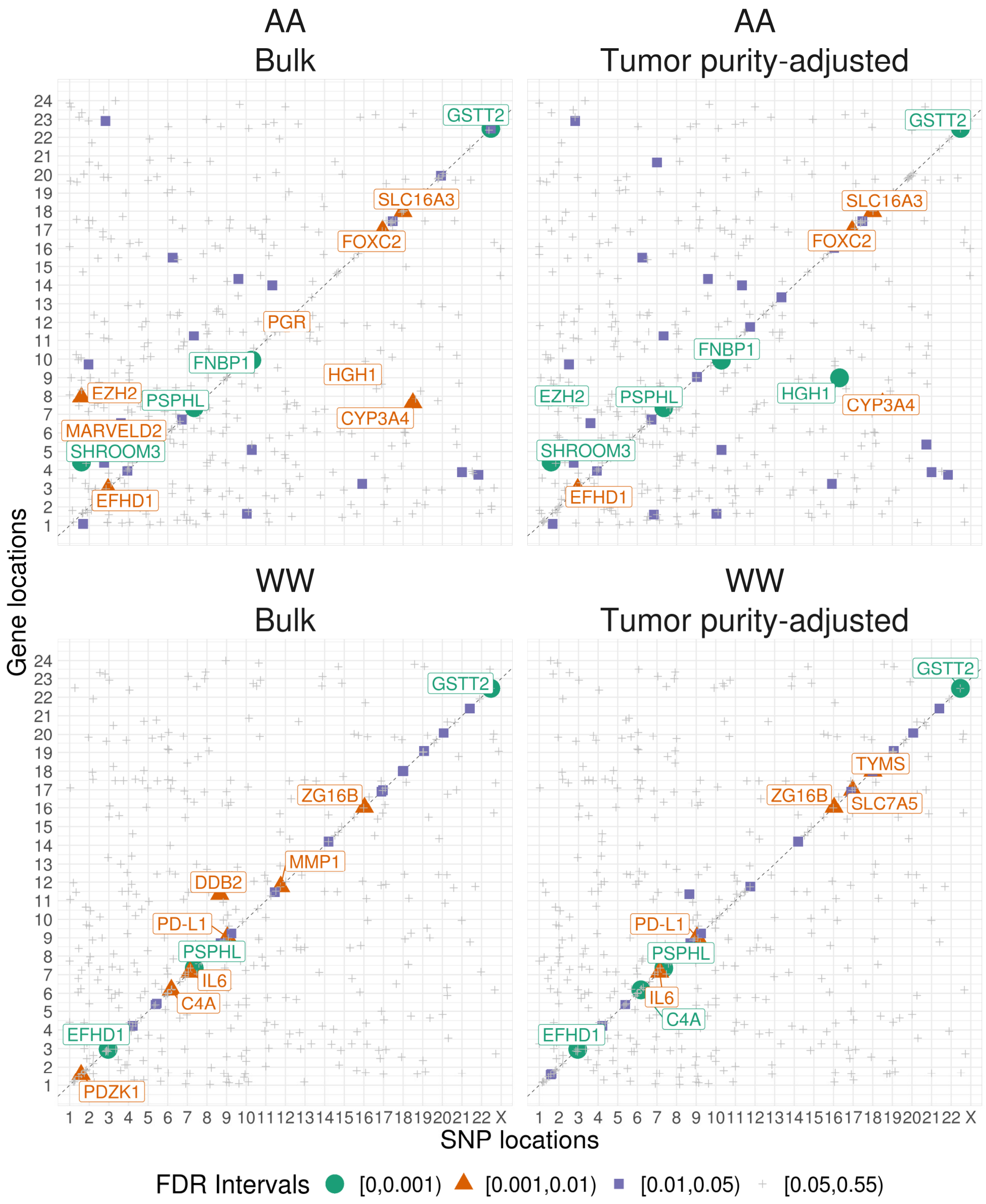


**Figure 4**: Cis-trans plots, as in Supplementary Figure 2, across self-identified race (top to bottom) and across adjustment for tumor purity (eQTLs in bulk tumor expression on left and eQTLs in tumor purity-adjusted expression on left)

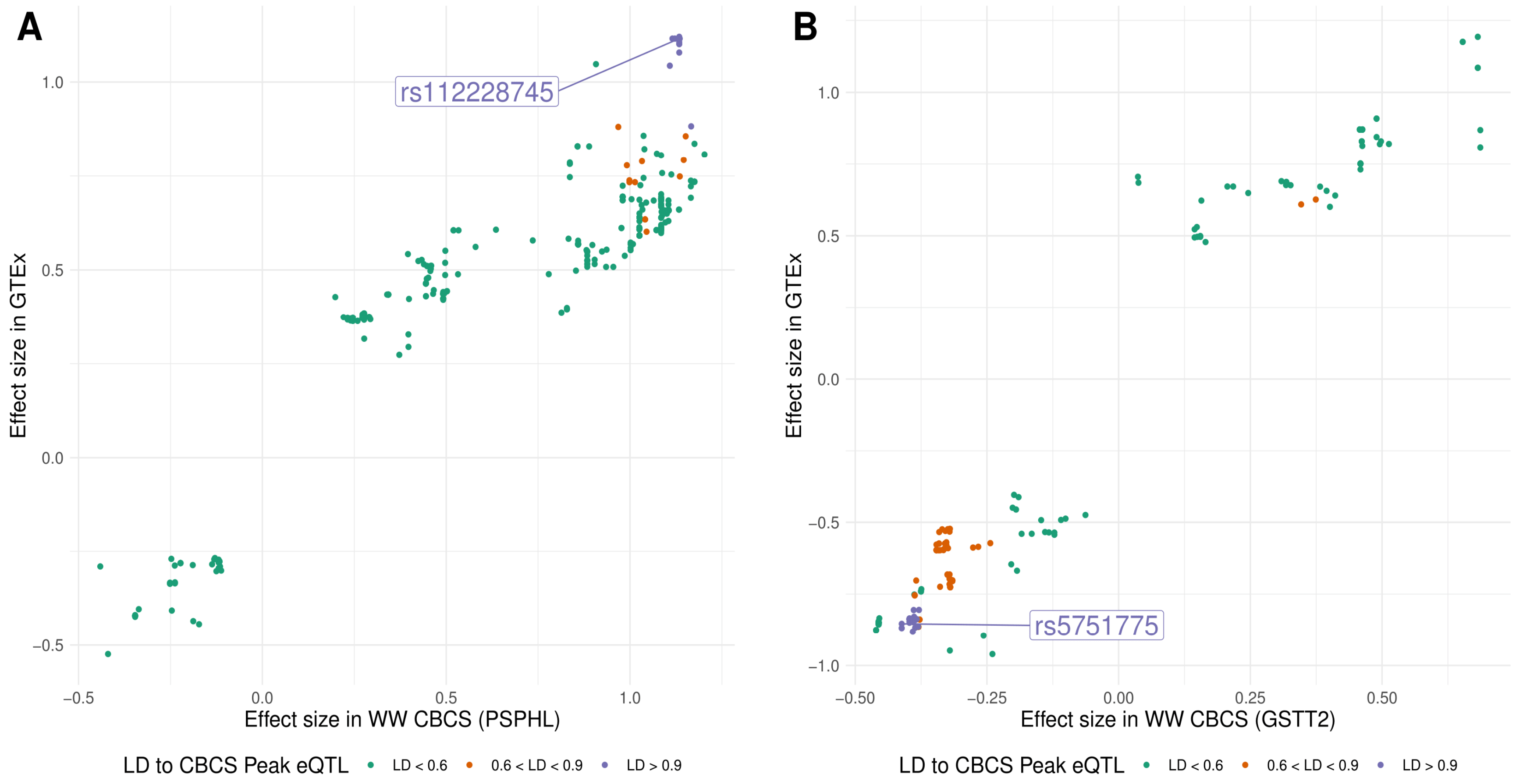


**Figure 5**: Each point represents a significant eQTL for *PSPHL* (A) and *GSTT2* (B) found in both GTEx and the CBCS WW sample, colored by the strength of linkage disequilibrium to the top eSNP in CBCS. Absolute effect size of significant eQTLs in WW CBCS is plotted on the X-axis and absolute effect size of significant eQTLs in GTEx multiplied by the sign of the effect size in CBCS is plotted on the Y-axis.

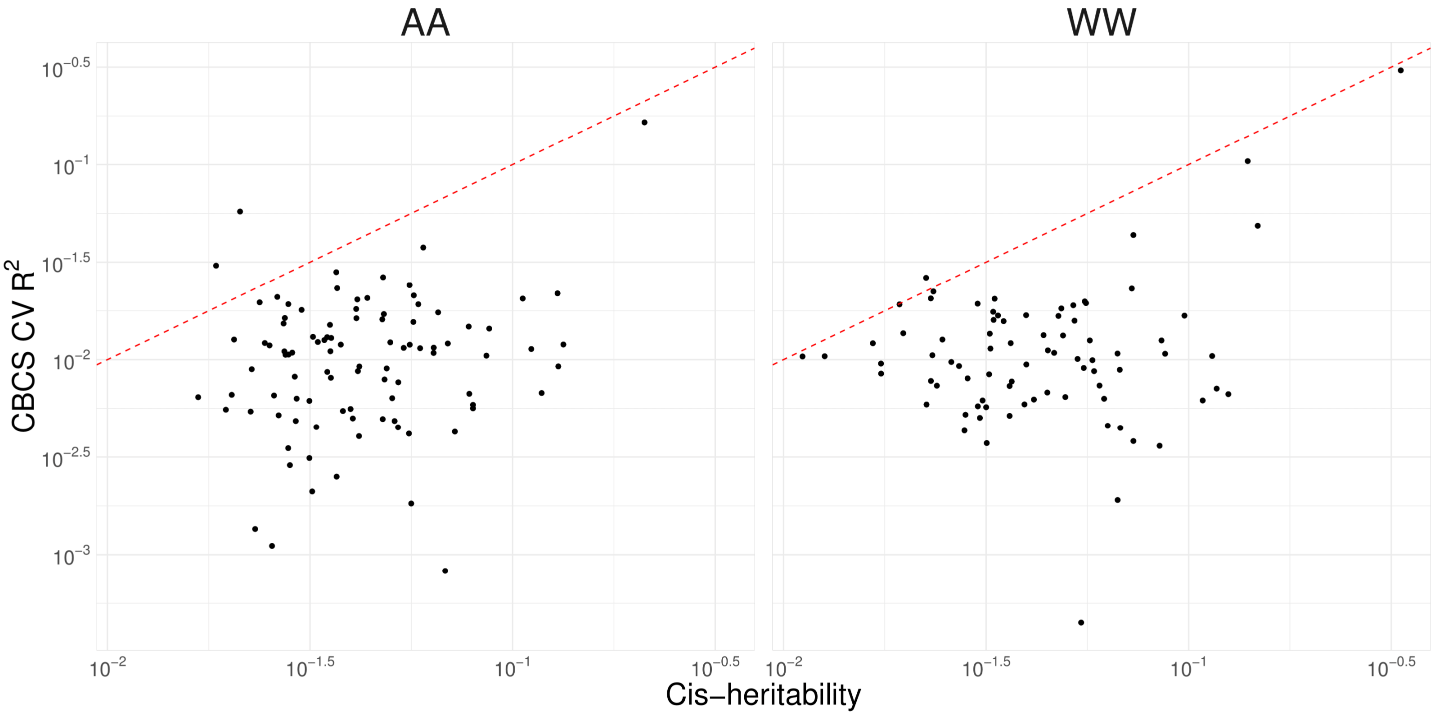


**Figure 6**: Comparison of cis-$h^2$ estimates (X-axis) and cross-validation $R^2$ (Y-axis) for each gene with likelihood ratio test $P < 0.10$ for cis-$h^2 = 0$ across AA and WW women in CBCS training set. The 45-degree line (i.e. $Y = X$) is provided for reference in red.

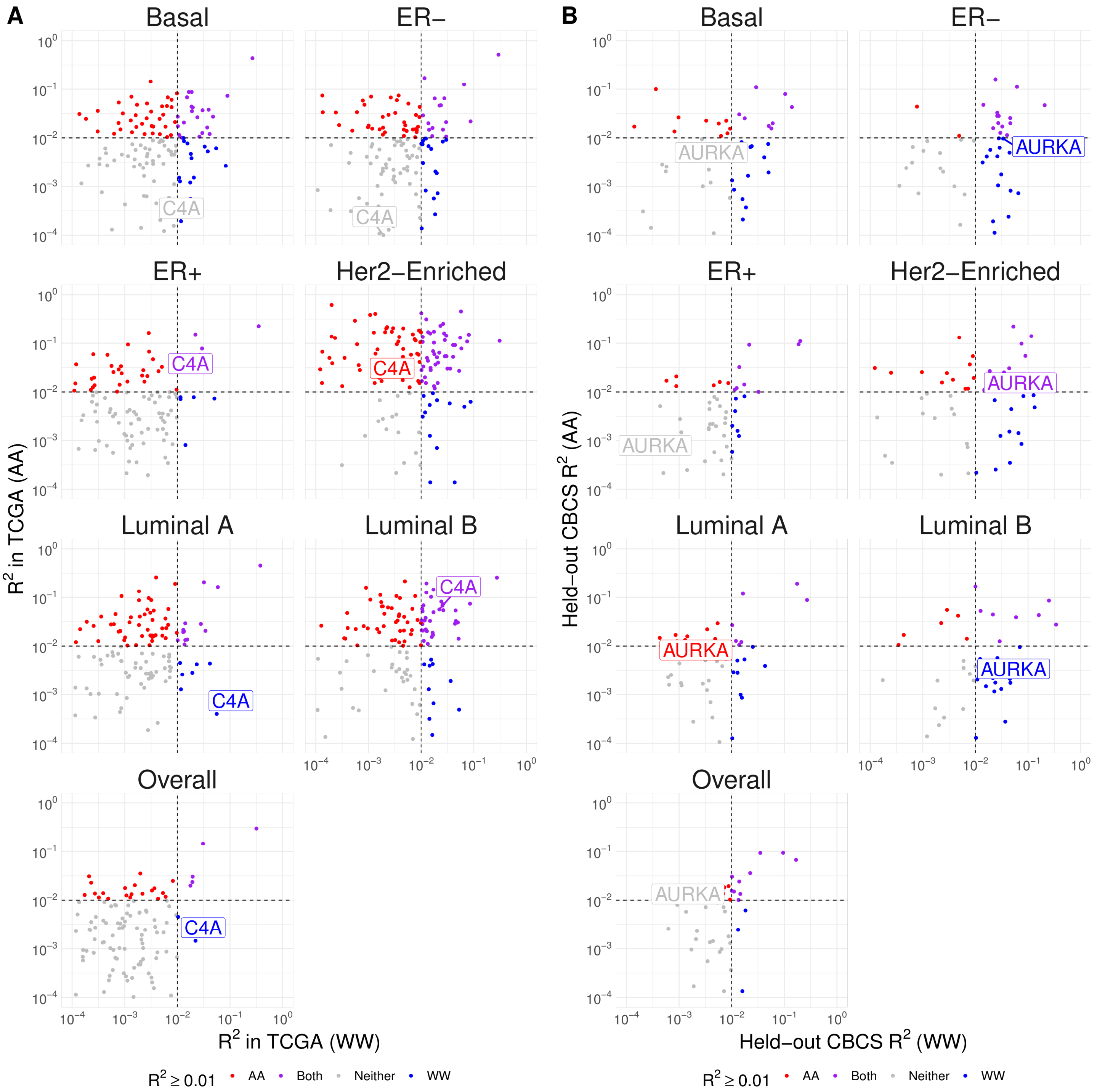


**Figure 7**: Comparison of prediction $R^2$ across race, stratified by PAM50 molecular subtype and estrogen receptor status in TCGA (A) and CBCS (B). Squared Spearman correlation in WW (X-axis) and AA (Y-axis) for each of the available genes are plotted. Note that both scales are logarithmic. Dotted lines represent $R^2 = 0.01$. Colors represent the model with which a given gene can be predicted at cross-validation $R^2 > 0.01$. A representative gene with variable $R^2$ across subtypes is labelled.

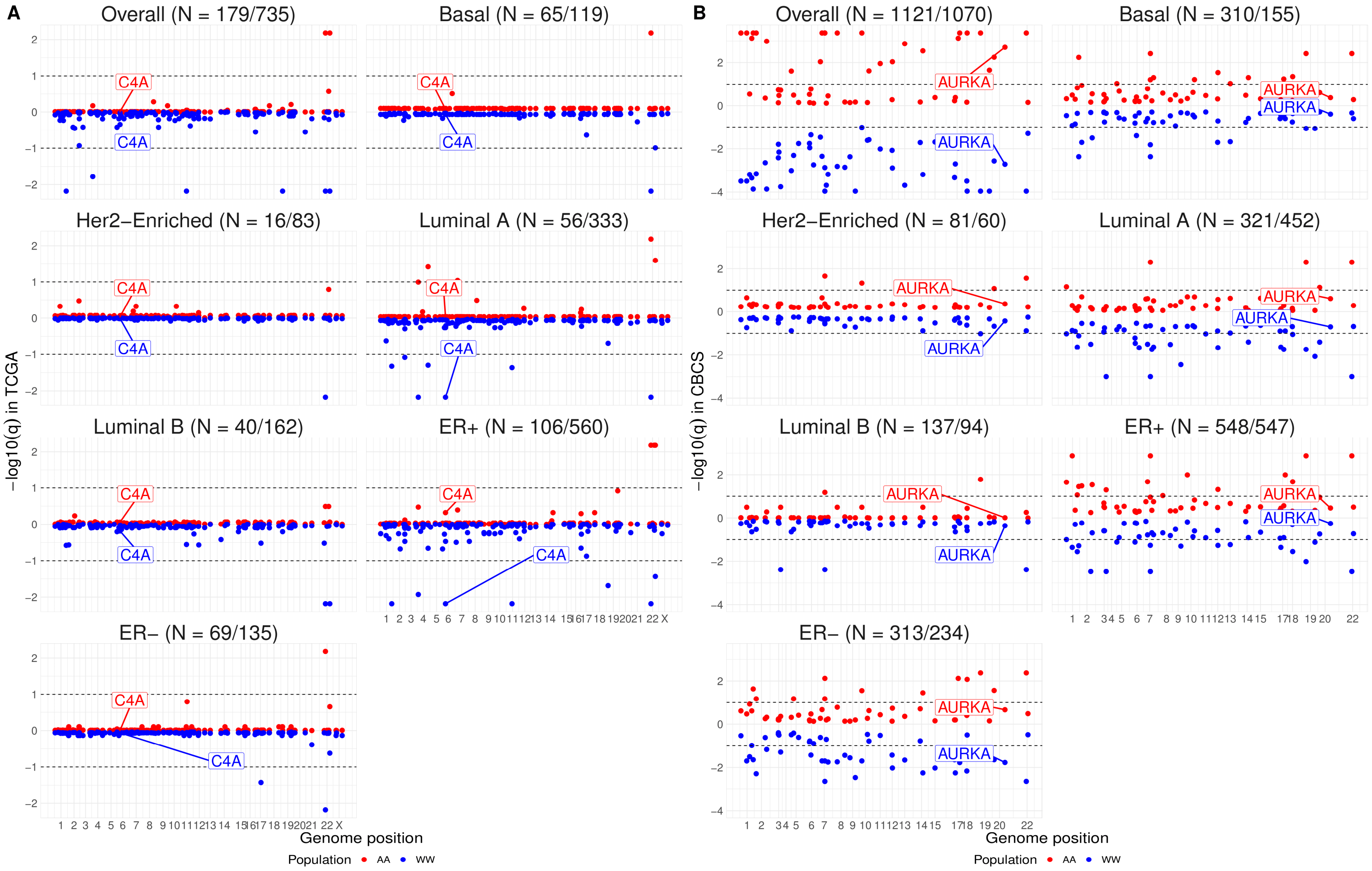


**Figure 8**: Storey's − log10 q-values from P-values of permutation tests over 10,000 permutations to assess significance of external validation $R^2$ in TCGA (A) and held-out CBCS (B). Dotted lines represent q = 0.10. Sample sizes are provided in the form (AA/WW). A representative gene with variable permutation q-value across subtype is labelled.

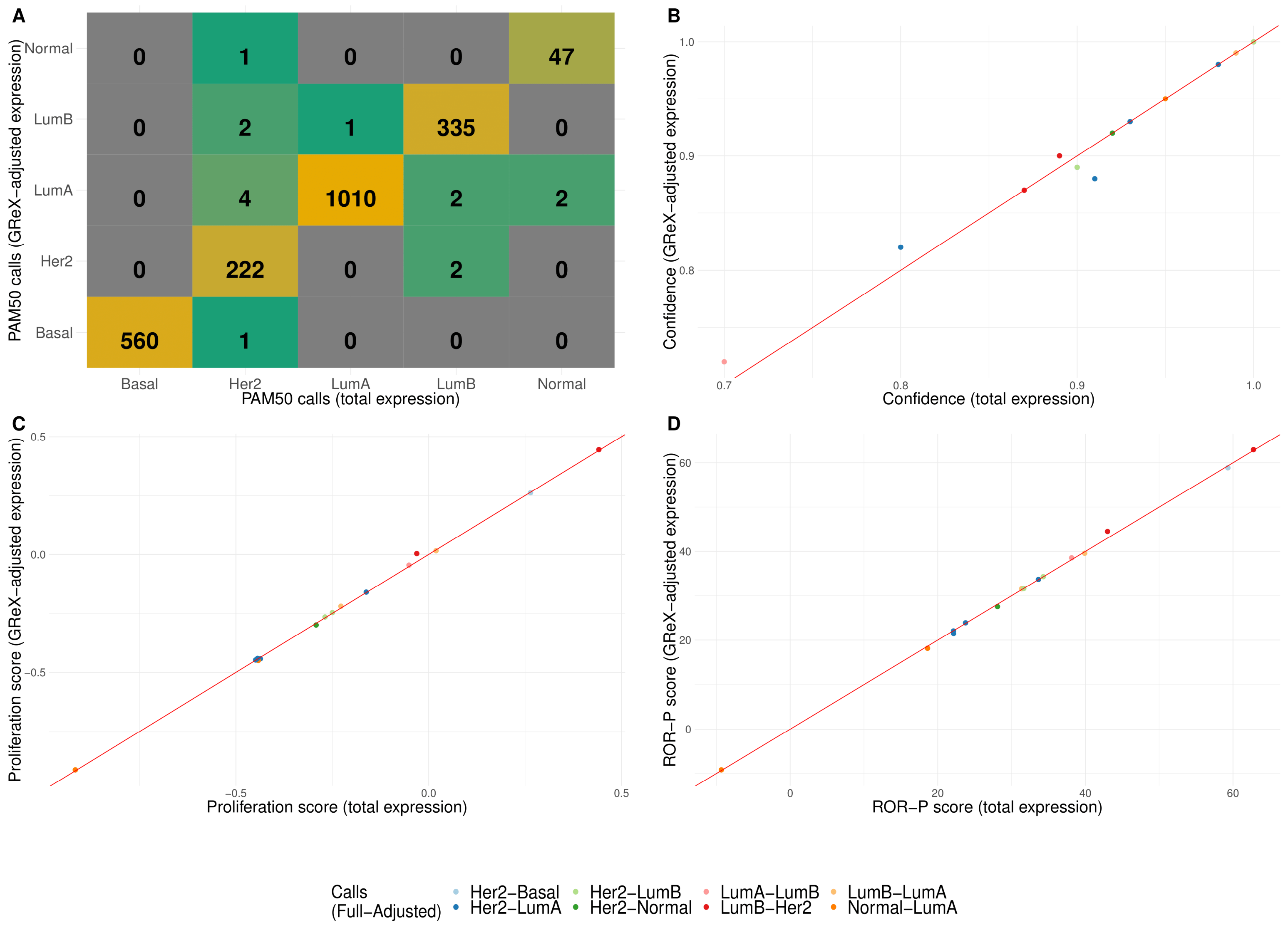


**Figure 9**: (A) Confusion matrix of PAM50 molecular subtype calls in held-out CBCS derived from full, unadjusted tumor expression(X-axis) and tumor expression adjusted for GReX (Y-axis) (B,C,D) PAM50 confidence (B), proliferation (C), and ROR-P (D) scores across adjustment for GReX. Points represent discordant calls across adjustment, colored by calls before and after adjustment. Red line provides the 45-degree regression line for reference. PAM50 confidence is defined as $1 - P$-value of a sample's Spearman correlation test to the PAM50 centroid. The proliferation and ROR-P scores are linear combinations of a sample's distances to all five PAM50 centroids and are measures of a tumor's proliferation and risk of relapse.

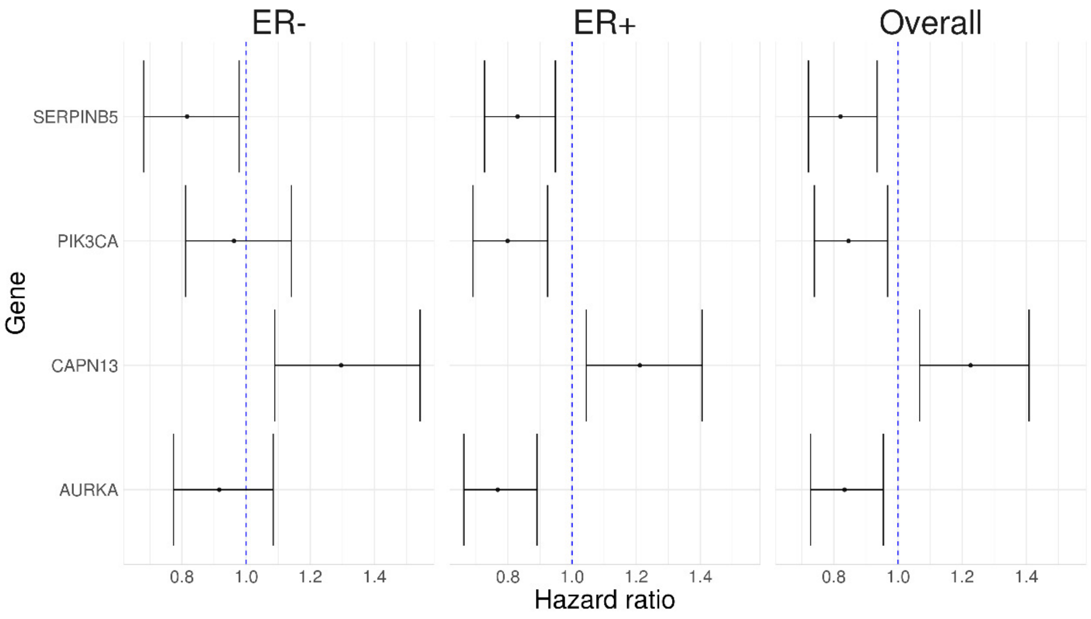


**Figure 10**: Caterpillar plots for hazard ratio of breast cancer-specific survival in AA women for an increase of one standard deviation of GReX across models unadjusted for estrogen receptor subtype and stratifying for estrogen receptor subtype.

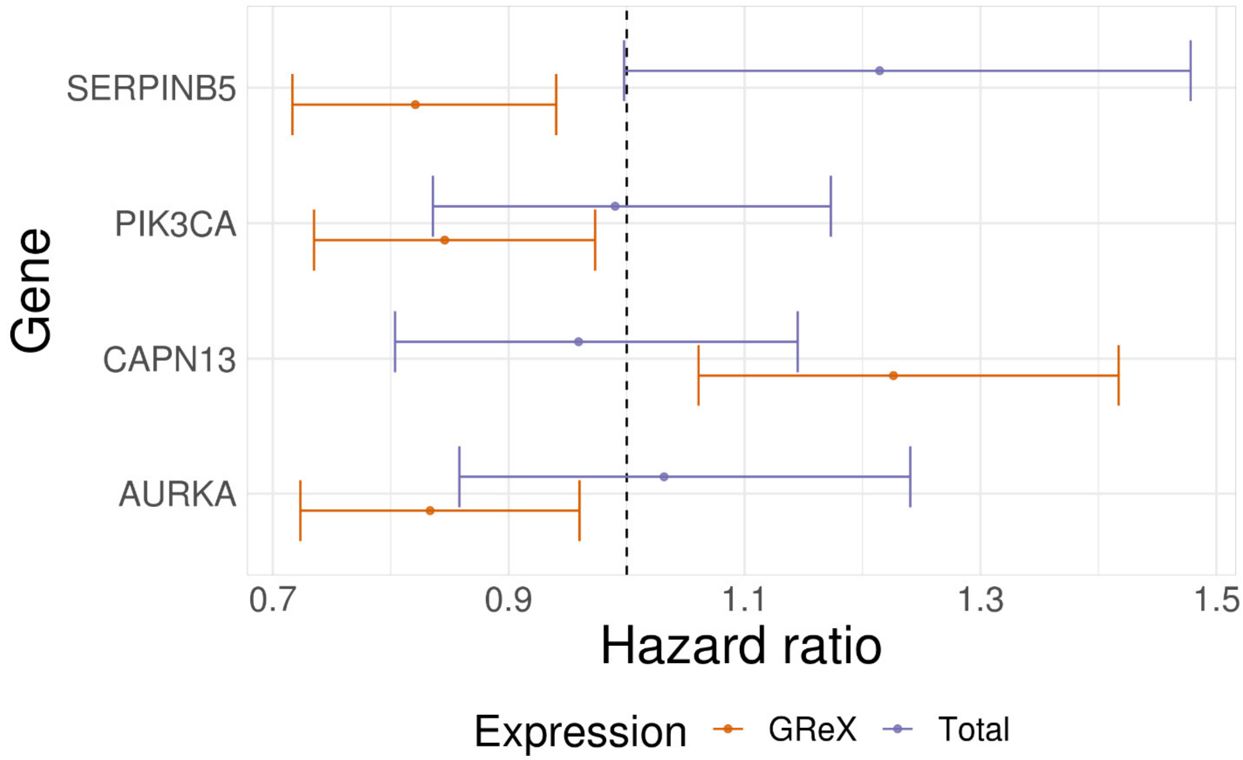


**Figure 11**: Hazard ratios and 95% confidence intervals, adjusted for false discovery via Benjamini-Hochberg, as estimated from breast cancer-specific Cox models in AA women. Association with total expression (purple) and GReX (orangle) of 4 TWAS-detected genes are compared.

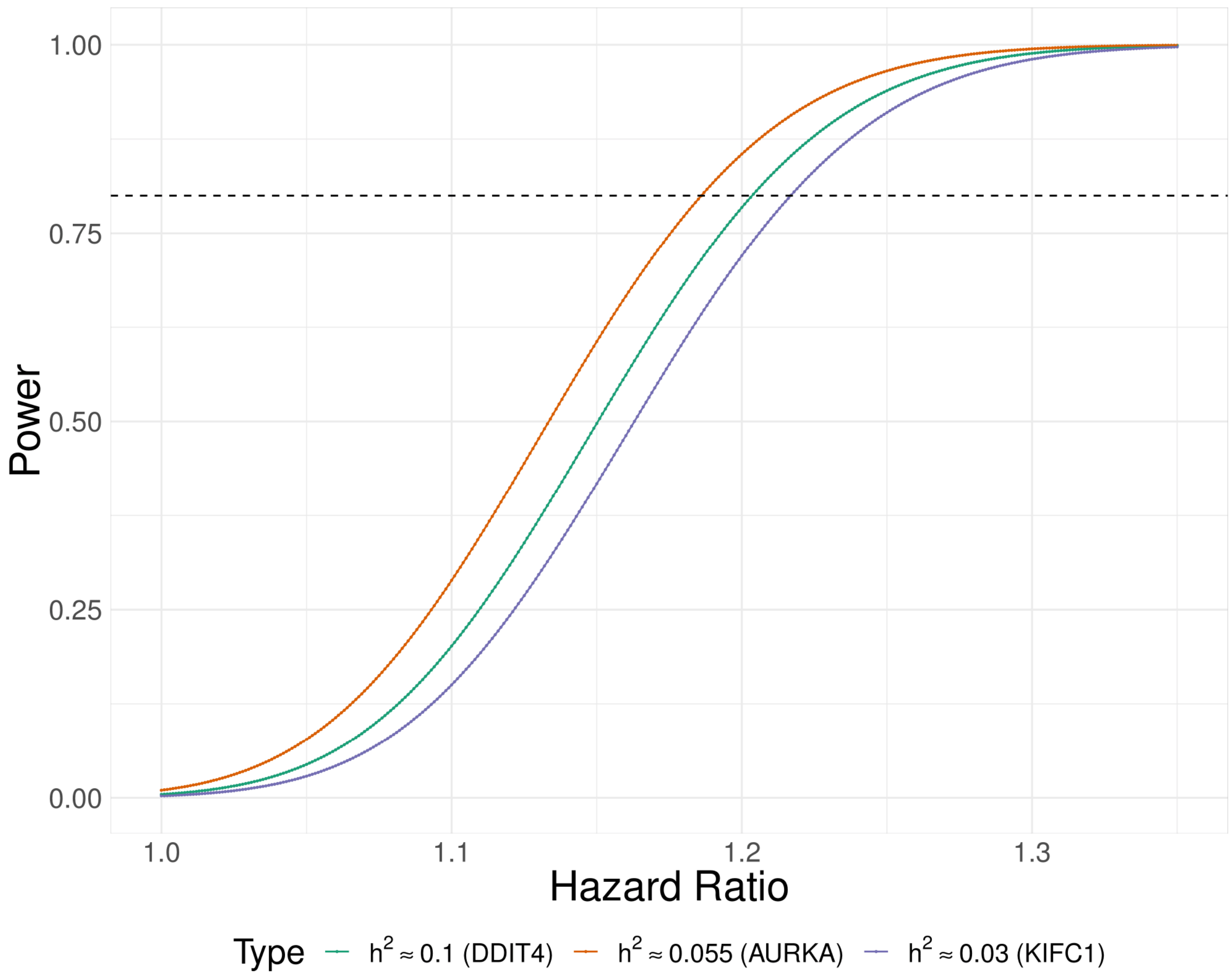


**Figure 12**: Comparison of power of TWAS in CBCS sample of $N = 3{,}828$ and 348 breast cancer-specific deaths. Power (Y-axis) to detect a given hazard ratio (X-axis) is plotted. Curves correspond to genes of varying cis-$h^2$: *DDIT4* has high $h^2$ across AA and WW, *AURKA* has average $h^2$ across AA and WW, and *KIFC1* has the lowest $h^2$ across AA and WW. Power calculations are derived from 1,000 re-samplings of the empirical distribution function of the GReX of of a given gene. Dotted line represents 80% power.

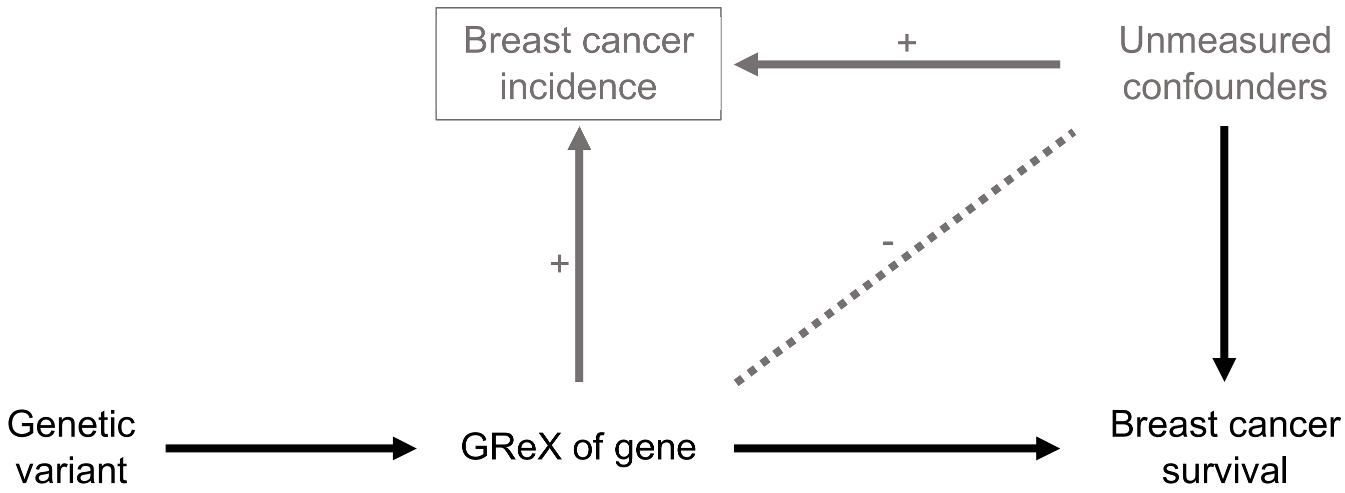


**Figure 13**: Modified from Paternoster et al. Directed acyclic graph that shows how collider bias is introduced (grey path) in case-only studies. Here, in this case-only study, we condition on breast cancer incidence, which may open up a potential collider bias with unmeasured confounders in the measure of association between the GReX of a gene and breast cancer survival.
